# Noncanonical circRNA biogenesis driven by alpha and gamma herpesviruses

**DOI:** 10.1101/2022.07.08.499347

**Authors:** Sarah E. Dremel, Vishal N. Koparde, Jesse H. Arbuckle, Chad H. Hogan, Thomas M. Kristie, Laurie T. Krug, Nicholas K Conrad, Joseph M. Ziegelbauer

## Abstract

Herpesviruses rely on the host transcriptional machinery for expression of their >100 different transcripts. To prioritize viral transcripts, herpesviruses affect significant changes in all stages of gene expression. Herein, we examined how herpesviruses alter circular RNA (circRNA) synthesis resulting in distinct characteristics for those expressed from the host or viral genome. CircRNA were identified with our Circrnas in Host And viRuses anaLysis pIpEline (CHARLIE) capable of *de novo* annotation using aggregated calls from five distinct tools. Comparative profiling for Herpes Simplex Virus-1, Kaposi sarcoma-associated herpesvirus, and murine gammaherpesvirus 68 identified thousands of back splicing variants, including circRNA species commonly expressed in all phases of infection. CircRNA expression was highest during lytic infection, with a transcriptional density 500-fold greater than the host. We characterized cis- and trans-elements controlling back splicing and found viral circRNAs are generally resistant to spliceosome inhibition or depletion, with >90% lacking canonical splice donor-acceptor sites. Using eCLIP and 4sU-Sequencing, we determined that the viral RNA binding protein, ORF57, enhanced synthesis for a subset of viral and host circRNAs. Our work elucidates a unique splicing mechanism driven by late lytic replication and identifies a class of transcripts with potential to function in replication, persistence, or tumorigenesis.

## INTRODUCTION

Herpesviridae is an extensive virus family with host-adapted species for a range of organisms including mollusks, fish, birds, reptiles, and mammals. There are nine herpesvirus species known to infect humans, including the alphaherpesvirus Herpes Simplex Virus-1 (HSV-1, human herpesvirus 1) and the gammaherpesvirus Kaposi sarcoma-associated herpesvirus (KSHV, human herpesvirus 8). HSV-1 commonly causes recurrent oral and genital lesions. The virus is also responsible for more severe disease including herpes keratitis, neonatal mortality and congenital malformations, and viral encephalitis^1–3^. Individuals are usually infected with HSV-1 during childhood, and by age 10 approximately 65% of the global population tests seropositive^4^. Infection with HSV increases the probability of HIV transmission and HSV reactivation is linked to increased HIV load^5,6^. KSHV is the etiological agent of several cancers, including Kaposi sarcoma and primary effusion lymphoma^7^. Additional KSHV-associated pathologies include Multicentric Castleman disease (MCD) and KSHV inflammatory cytokine syndrome (KICS). KSHV seroprevalence varies by region and is highest in sub-Saharan Africa (>50%), the Mediterranean (20-30%), and among individuals living with HIV/AIDS in the U.S (∼40%)^7,8^. A closely related virus, murine gammaherpesvirus 68 (MHV68) or murid herpesvirus-4 (MuHV-4), has high genetic homology and serves as a tractable animal model for chronic gammaherpesvirus infection^9,10^. Despite decades of study, a therapeutic agent capable of clearing these viruses is lacking. Of the nine human herpesviruses, only varicella zoster virus has an FDA-approved vaccine.

Herpesviruses have large double-stranded DNA genomes and establish latent reservoirs that persist for the life of the host. These viruses employ a biphasic life cycle, which includes a lytic (replicative) phase and a latent (quiescent, immune evasive) phase. To facilitate their complex, tissue-specific life cycle, herpesviruses express upwards of 100 different transcripts. These include messenger RNAs (mRNAs), long noncoding RNAs (lncRNAs), microRNAs (miRNAs), and a recently discovered class of transcripts— circular RNAs (circRNA). CircRNAs are single-stranded RNA species formed by a 5’ to 3’ covalent linkage. The mechanism underlying circRNA synthesis, back splicing, is catalyzed by the spliceosome and promoted by RNA binding proteins (RBPs) or tandem repeat elements which mediate interaction of back splice junction (BSJ) flanking sequences^11–15^. CircRNAs are an alternative splicing product and thus expand the number of transcripts without extending genome size; a strategy that may be advantageous for organisms with limited coding space.

In 2018, three groups published discovery of virus-derived circRNA^16–18^. Viral circRNAs have now been experimentally confirmed for beta- and gamma-herpesviruses, human papillomavirus, Merkel cell polyomavirus, hepatitis B virus, respiratory syncytial virus, and coronaviruses^16–26^. In addition, bioinformatic evidence supports expression of circRNAs from other viral genomes^27^. Host circRNAs have been shown to function as miRNA sponges, protein scaffolds or sponges, transcriptional enhancers, and translational templates^28^. Less is known regarding the function of viral circRNAs; some serve as translational templates and are packaged in virions and extracellular vesicles^22,23^. Gammaherpesvirus circRNAs have been shown to modulate cell growth and apoptosis, pathways of particular interest for oncogenic viruses^17,29^. CircRNAs have also been proposed to be putative biomarkers for cancer, with cancer-specific expression profiles and a global anti-correlation between circRNA steady-state levels and cell proliferation^30,31^. In line with this, our lab detected KSHV circRNAs in lymph node biopsies obtained from primary effusion lymphoma patients^29^. CircRNAs are currently being investigated as a mechanism for mRNA-vaccine delivery as they have low immunogenicity, long half-lives, and the potential for cap-independent translation^32–34^.

As alternative splicing products, circRNA share almost complete sequence complementarity with their linear counterpart derived from the same gene. The only unique sequence is their 5’ to 3’ BSJ. Research into circRNAs has increased exponentially with the advent of high throughput sequencing (HTS). HTS paired with chimeric transcript analysis now enables global circRNA detection and quantification^35,36^. These techniques have found that circRNAs are ubiquitously expressed in an array of organisms and tissues^37,38^. Herein, we characterize the circRNAome of the human herpesviruses HSV-1 and KSHV, as well as that of MHV68, a mouse model for KSHV. We used cell culture and mouse models to thoroughly profile circRNAs expressed during lytic, latent, and reactivation phases. A custom bioinformatic pipeline, called CHARLIE (Circrnas in Host And viRuses anaLysis pIpEline) was developed to facilitate high-throughput analysis of viral circRNAs. Using these approaches, we identified thousands of viral circRNAs, some of which approach abundance of the housekeeping gene *GAPDH*. Notably, we are the first to identify HSV-1 encoded circRNAs, including circular transcripts derived from the middle exon of ICP0 and the stable lariat intron of the latency-associated transcript (LAT). Next, we characterized cis- and trans-acting factors which promote circRNA biogenesis. We found that most viral circRNA species were resistant to major spliceosome inhibition and lacked canonical splice donor-acceptor sites. Finally, using eCLIP and Nascent (4sU) RNA-Seq, we determined that the KSHV RNA binding protein (ORF57) enhanced circRNA synthesis for a subset of viral and host transcripts. Our work identifies dozens of novel herpesvirus transcripts and elucidates a unique splicing mechanism driven by lytic infection.

## RESULTS

### Herpesvirus circRNA repertoire

We used models for primary lytic infection, latency, and reactivation to explore circRNAs expressed during these distinct transcriptional programs (Fig. 1A, Supplementary Figure 1-1). HSV-1 models include fibroblasts (MRC-5) infected for 12 or 24 hours, murine trigeminal ganglia (TG) latently infected for four weeks, and murine TG from latently infected animals explanted (TG explant) for 12 hours with or without a reactivation enhancer (JQ1). KSHV models include human dermal lymphatic endothelial cells (LEC) or human umbilical vascular endothelial cells (VEC) infected for three days, and iSLK-BAC16 treated with doxycycline (Dox) and sodium butyrate (NaB) for three days. MHV68 models include mouse fibroblasts (3T3) infected for 18 hours, latently infected splenic germinal center B-cells (GC B-cell) from mice infected for 16 days, and A20 HE-RIT (HE-RIT) treated with Dox and tetradecanoyl phorbol acetate (TPA) for 24 hours. External RNA Controls Consortium (ERCC) synthetic spike-ins were added to total RNA, which was ribosomal RNA (rRNA) depleted and sequenced to generate 150 base pair (bp) paired-end reads. If indicated, total RNA was digested with ribonuclease R (RNase R) to enrich circRNAs prior to sequencing.

**Fig. 1.**
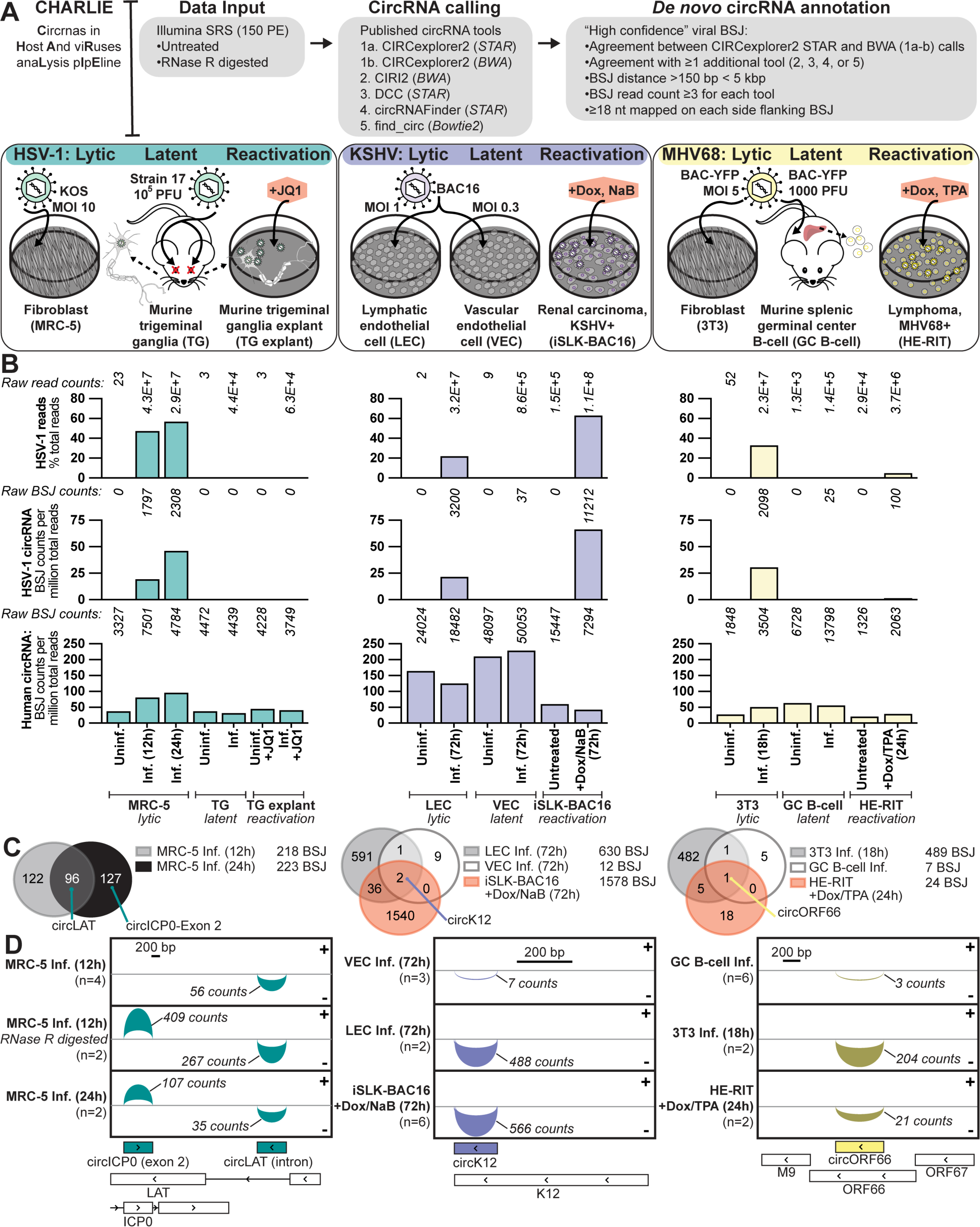
Herpesvirus circRNA repertoire. High confidence circRNAs were called from 150 paired end (PE) Illumina short read sequencing (SRS) data using. Infection models include HSV-1: infected fibroblasts (MRC-5), latently infected murine trigeminal ganglia (TG), and latently infected murine TG explanted (TG explant) with or without a reactivation enhancer (JQ1); KSHV: infected human dermal lymphatic endothelial cells (LEC) or human umbilical vascular endothelial cells (VEC), iSLK-BAC16 reactivated with doxycycline (Dox) and sodium butyrate (NaB); MHV68: infected mouse fibroblasts (3T3), latently infected murine germinal center B-cells (GC B-cell), and A20 HE-RIT (HE-RIT) reactivated with Dox and tetradecanoyl phorbol acetate (TPA). B) All reads mapped to the viral genome as a percentage of total reads. The sum of raw counts for each condition are reported above. High confidence viral or host circRNAs plotted as BSJ counts per million total reads. The sum of raw BSJ counts for each condition are reported above. Column bars are the average of biological replicates. C) Venn Diagrams of overlapping high-confidence viral back splice junction (BSJ) variants. D) Sashimi plots for select high confidence circRNA variants, arcs are proportional to raw BSJ counts.

CircRNAs were identified by their unique sequence element, the back splice junction (BSJ), a 5’ to 3’ covalent linkage that generates the continuous loop. Our custom pipeline, CHARLIE (Circrnas in Host And viRuses anaLysis pIpEline), aggregates BSJ calls from CIRCexplorer2^39^, CIRI2^40^, DCC^41^, circRNAFinder^42^, and find_circ^43^ (Fig. 1A). BSJ calls must meet the following criteria: ≥3 counts, 18 nucleotides (nt) mapped on both sides of the BSJ, circRNA splice donor acceptor distance ≥150 nt, viral circRNA splice donor acceptor distance ≤5000 nt. High confidence circRNAs were those in agreement between CIRCexplorer2 STAR^44^ and BWA^45^ mapping outputs and called in at least one other tool. These criteria were in line with recent recommendations^46,47^ for best circRNA practices, requiring agreement between at least two *de novo* circRNA annotation tools reliant on distinct RNA-Seq mapping programs. Importantly, this approach enables *de novo* circRNA calling for genome assemblies lacking curated transcript features, e.g. transcription start sites, transcription end sites, exons, introns, as is the case for viral genomes. Using this approach, we identified thousands of high confidence BSJs with significant overlap between biological replicates (Supplementary Figure 1-2). The degree of overlap is likely an underestimate as we required exact matching and would likely increase if we allowed +/- 5 nt of wobble for each BSJ position. The median distance between back splice donor and acceptor was 334, 447, and 907 nt for HSV-1, KSHV, and MHV68 circRNAs, respectively (Supplementary Figure 1-3). The median distance between back splice donor and acceptor for host circRNAs was ∼7,000 nt. The discrepancy in BSJ distance is expected in the context of average gene size; 1-4 kb and 10-15 kb for viral and host genes respectively. Our findings are summarized in a resource table with the genomic position, BSJ flanking sequence, splice donor-acceptor, and raw read count per condition for all high confidence viral BSJs (Supporting Dataset 1). Overall mapping statistics and circRNA calls are reported in Supplementary Tables 1 and 2.

To validate our sequencing results, we performed RNase R resistance assays. In this assay, linear molecules with 5’ and 3’ ends are substrates for the endonuclease and degraded, while circularized molecules are resistant to digestion. We assessed RNase R digested material using RNA-Seq (Sup Fig. 1-4A) and divergent qPCR (Sup Fig. 1-4B). As expected, RNase R digestion resulted in an enrichment of BSJ reads detected (Supplementary Tables 2). There was significant overlap between untreated and RNase R treated samples with 102 out of 218 HSV-1 BSJ (47%) and 80 out of 630 KSHV BSJ (13%) in common (Sup Fig. 1-4A). In the same samples 311 out of 924 (34%) and 1358 out of 2664 (51%) human BSJ were in common between untreated and RNase R treated samples. Using divergent primer qPCR amplification, we found all herpesvirus circRNAs tested were either resistant or enriched after RNase R digestion (Sup Fig. 1-4B).

Viral circRNA abundance echoed total read composition, with the highest number detected during lytic infection (Supplementary Tables 1 and 2, Figure 1B). Host circRNA expression was less dynamic and largely influenced by cell type; with primary endothelial cells (LEC, VEC) possessing the most BSJ reads (Figure 1B). During primary lytic infection, 218, 630, and 489 unique BSJ variants were expressed by HSV-1, KSHV, and MHV68, respectively (Figure 1C). As a function of genome size, herpesviruses expressed >1,000 BSJ variants per megabase pair (Supplementary Table 2). By comparison, the host expressed ∼7 BSJ variants per megabase pair. To make this calculation, we used a conservative estimate that 100% of the viral genome was transcribed and 10% of the host genome was transcribed. Based on these values, the circRNA transcriptional density of the herpesvirus genomes is strikingly high—at least 500-fold greater than the host.

We compared the repertoire of viral BSJ variants expressed in our models. Notably, KSHV circK12 and MHV68 circORF66 were expressed in lytic, latent, and reactivation models (Figure 1C-D). No HSV-1 BSJ variants were detected in murine models with our standard cutoff of ≥3 BSJ reads (Figure 1C, Supplementary Table 2). As viral gene expression is particularly restricted during latency (Supplementary Figure 1-1), we lowered our cutoff to ≥1 BSJ reads and reanalyzed high confidence circRNAs. With the lower threshold, we identified a dozen BSJ variants derived from the HSV-1 latency associated transcript (LAT) (Supplementary Figure 1-5). Of these, circLAT_KT899744.1: 6137-6814 was expressed in murine trigeminal ganglia explants (latent) and infected human fibroblasts (lytic). circLAT was also validated with RNase R assays in latently infected murine TG (Supplementary Figure 1-4B), suggesting that some viral circRNAs may be filtered out with the stringent criteria used in Figure 1. One biological function of circRNAs is to serve as miRNA sponges^28,48^. In this way circRNAs modulate miRNA-mediated mRNA depletion. We used the sequence identity of HSV-1 circLAT, KSHV circK12, and MHV68 circORF66 to predict circRNA-miRNA, and miRNA-mRNA interaction partners (Supplementary Fig. 1-6A-C). Downstream phenotypes that may ultimately be controlled by these circRNA-miRNA-mRNA interaction networks included senescence and mTOR signaling (Supplementary Fig. 1-6D). Our profiling identifies novel viral transcripts with the potential to influence herpesvirus pathogenesis during all phases of infection.

### Profile of high-incidence lytic circRNAs

Fig. 2 includes all high confidence circRNAs detected in primary lytic models. Sashimi plots were limited to circRNA events found in at least two biological sequencing replicates. Sequencing reads were segregated to visualize traces containing BSJs, “circ”, versus nonchimeric or “linear” transcripts (Fig. 2A). Viral circRNA traces mimicked that of linear transcripts, with select genes being hotspots (>50 raw BSJ reads) for back splicing (Figure 2). These included circRNAs colinear with HSV-1 ICP0, LAT, UL12, UL19, UL36, and UL42; KSHV PAN, vIRF4, and K12; and MHV68 ORF66 (Fig. 2B-D, Supporting Dataset 1). During HSV-1 lytic infection, we detected circularization of the middle exon of ICP0 and a truncated version of the LAT lariat intron (Supplementary Figure 2-1A). There was also a highly abundant circRNA derived from the HSV-1 processivity factor, UL42 (Supplementary Figure 2-1A). Consistent with prior profiling^17,18,20^, major circRNA species expressed by KSHV include a BSJ variant within vIRF4 and a BSJ cluster in PAN (Figure 2C, Supplementary Figure 2-1B). We also identified an additional high incidence species, colinear to KSHV K12. Four BSJ variants comprised this cluster, with a single 3’ splice acceptor (NC_009333.1: 117534) and slightly variable 5’ donor (NC_009333.1: 117686, 117687, 117688, 117689). Low abundance circK12 species were previously detected in reactivated B-cell models^18^, however the species identified in our adherent cell models is distinct from the previously described circRNA. The major circRNA species expressed by MHV68 was a cluster within ORF66-69, with less abundant circRNAs expressed from ORF17 and ORF75A-C. The cluster within MHV68 ORF66-69 is consistent with findings from Ungerleider *et al.* (2018).

**Fig. 2.**
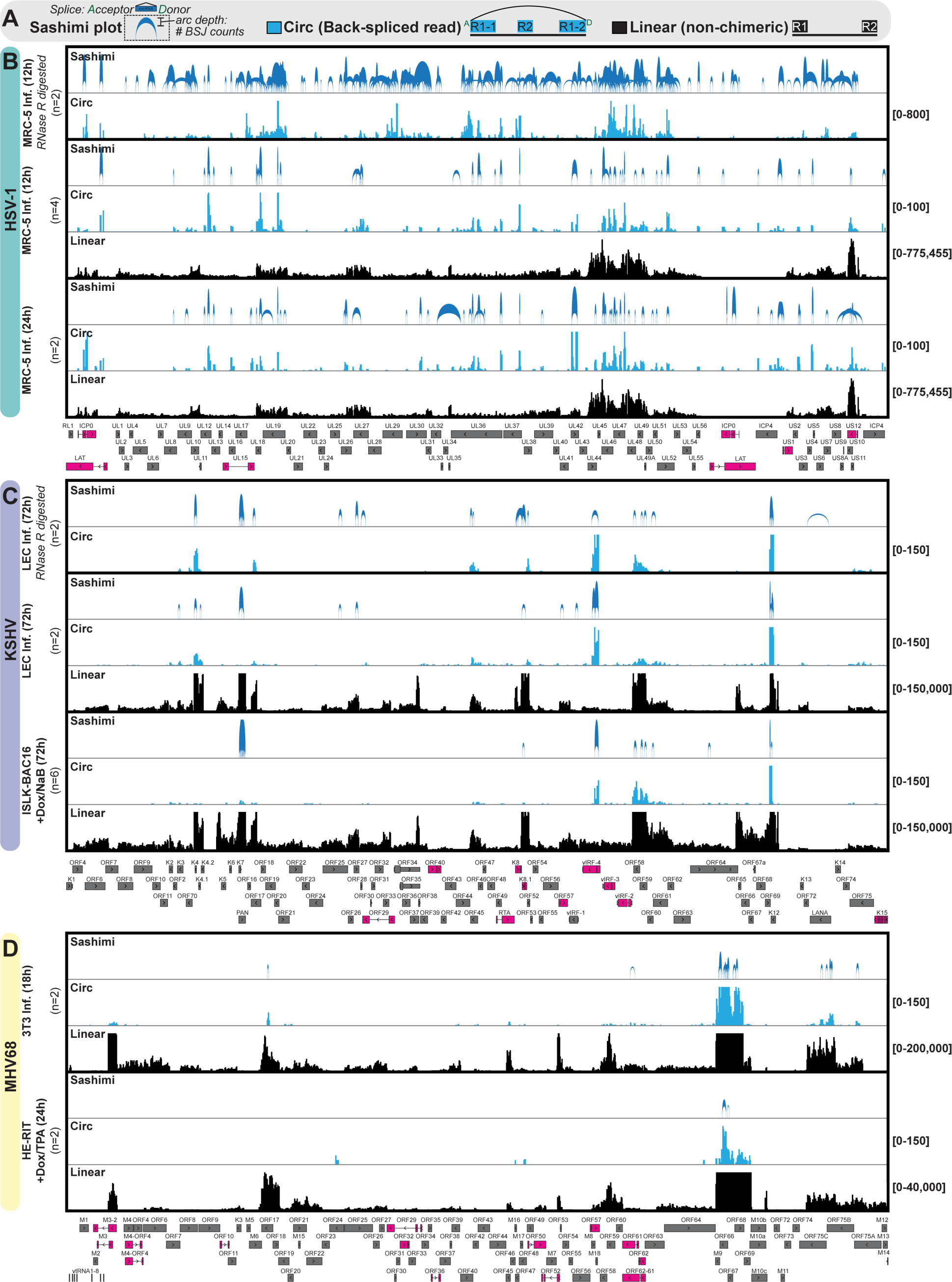
Profile of high-incidence lytic circRNAs. Visualization of high-confidence circRNAs for HSV-1, KSHV, and MHV68 in lytic models described in Fig. 1. If indicated, RNA samples were enriched for circRNAs using RNase R treatment (+RNaseR) prior to sequencing. Sashimi plots are limited to circRNA variants identified in at least two biological replicates, arcs are proportional to raw BSJ counts. Blue and black traces include circular (back spliced reads) and linear (non-chimeric) reads, respectively. Traces are the sum of raw BSJ or linear read values for all biological replicates. Y-axis minimum and maximum values are shown on the right. Single exon (grey) and multi-exon (pink) viral genes are shown below.

To determine copy levels of select high confidence circRNAs, we performed digital droplet PCR using divergent primers which only amplify circular transcripts (Supplementary Fig. 2-2). We included host transcripts, such as *7SK*, *GAPDH*, and *circHIPK3* for reference. Viral circRNAs ranged in abundance from 10^3^ to 10^6^ copies per ug total RNA. HSV-1 UL44 and KSHV PAN were the most abundant, reached 10^5^ and 10^6^ copies/ug total RNA, respectively. These levels are similar or greater than *circHIPK3*, a highly abundant human circRNA with published roles in miRNA sponging^49^. These findings identify back splicing hotspots within the viral genome, with select species reaching abundance only 100-fold less than the housekeeping transcript, *GAPDH*.

### Cis-elements coordinating viral back splicing events

When visualizing data mapped to the viral genome, we observed many circRNAs derived from single exon genes (Fig. 2). The lack of relationship between forward and back splicing was particularly notable for HSV-1, with circRNA species tiling the viral genome (Fig. 2B). This is perhaps unsurprising as only five of the ∼80 genes are spliced and HSV-1 itself inhibits the host splicing machinery^50,51^. Thus, we investigated flanking cis-elements to compare high confidence host and viral circRNA species (Fig 3A). First, we determined the 2 bases 5’ and 3’ to BSJ, as these would be the splice donor-acceptor (DA) sites. As an alternative splicing event, circRNAs are expected to rely on canonical splice DA signals, such as GT-AG^11,12^. As expected, 84-97% of host circRNAs used GT-AG as their splice-donor acceptor (Fig 3B, Supplementary Fig. 3-1A). Of the top ten most highly expressed species for each virus, two HSV-1 circRNA (circICP0-exon2, circUL29), two KSHV circRNA (circvIRF4, circPAN), and zero MHV68 circRNA possess GT-AG or CT-AC as their flanking splice DA (Supporting Dataset 1). Looking at all high confidence viral circRNAs, we found that only 0.6-7% used the canonical splice DA (Fig 3B). Similar trends were observed for circRNAs identified in KSHV and MHV68 reactivation models (Supplementary Fig. 3-1A). Analysis of circRNAs detected in RNase R digested samples corroborated our findings (Supplementary Fig. 3-1A). In summary >90% of viral circRNAs are not flanked by canonical splice DA elements, such as GT-AG or CT-AC.

**Fig. 3.**
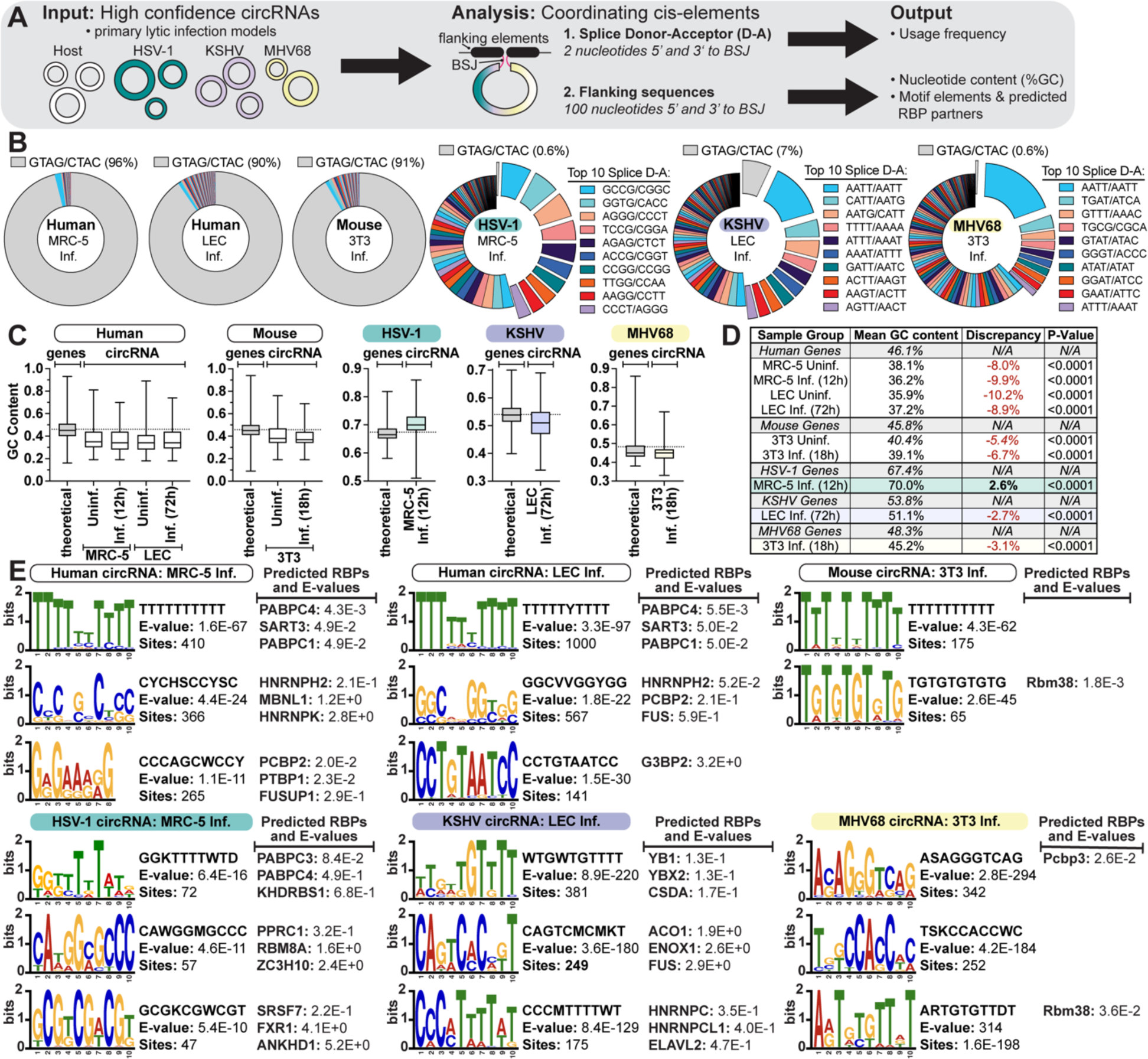
Cis-elements coordinating viral back splicing events. A) Infographic for BSJ flanking cis-element analysis performed on high-confidence viral and host circRNAs identified in primary lytic infection models. B) Splice donor-acceptor frequency for high confidence BSJ variants, reported as sense and antisense sequences. The percentage of the total which use the canonical splice donor acceptor (GT-AG/CT-AC) are reported above. C) GC content of 100 nucleotide (nt) BSJ flanking sequences relative to the theoretical GC content of genes for an organism. D) Wilcoxon t-tests were performed, relative to the theoretical gene GC content, to test significance. E) Nucleotide consensus plots, E-values, and number of sites in input sequences is given for BSJ flanking motifs. Motifs had p-values <0.05 and only the top three most significant (based on E-Value) are shown. Predicted RBP binding partners for BSJ flanking motifs. Predictions had p-values <0.05 and only the top three most significant (based on p-value) are shown.

Next, we evaluated GC content of the 100 nt flanking the BSJ (Fig 3C-D). GC content directly impacts many facets of gene expression including, transcription rates, splicing, and secondary structure. Previous work has found that low GC content is a significant predictor of circRNA hotspots^52^. Consistent with this, we found the GC content of external sequences flanking human and mouse circRNAs was significantly lower than the theoretical average for host genes (Fig 3C-D). This effect was most striking for human circRNAs with an average decrease of 9% relative to host gene GC content. GC content in KSHV and MHV68 circRNA flanking sequences followed a similar trend, with a decrease of 2.7 and 3.1%, respectively. Viral circRNAs from lytic reactivation models had more drastic decreases in GC-content with 7.4% for KSHV and 5.6% for MHV68 (Supplementary Fig. 3-1B). HSV-1 circRNAs were the outlier with a GC content increase of 2.6%; this increase was relative to the already high GC-content of HSV-1 genes (67%). To put this in context, HSV-1 circRNA flanking sequences had a GC content of 70% as compared to host derived circRNAs with a GC content of 36%. All of our findings were consistent when comparing circRNA identified in untreated or RNase R digested samples (Fig. 3C-D, Supplementary Figure. 3-1B).

To identify novel motifs with the potential to recruit RBPs and coordinate circRNA synthesis, we performed motif enrichment analysis. Supporting our findings in Fig. 3C, we found thymine tracks to be overrepresented in host circRNA flanking sequences (Fig. 3E). This was by far the most enriched motif with E-values of 1.6E-67, 3.3E-97, and 4.3E-62. Whether TTTTTTTTTT truly functions as a motif to coordinate protein binding or is merely indicative of the low GC content in host circRNA flanking regions remains to be explored. Distinct motifs were found for regions flanking viral circRNAs, with the most enriched being HSV-1: GGKTTTTWTD, CAWGGMGCCC, GCGKCGWCGT; KSHV: WTGWTGTTTT, CAGTCMCMKT, CCCMTTTWT; MHV68: ASAGGGTCAG, TSKCCACCWC, ARTGTGTTDT. Of these motifs, three contain thymine tracks, similar to host circRNA flanking elements. Five (ANKHD1, FUS, HNRNPH2, HNRNPK, PCBP2) of the 27 predicted RBP interaction partners have been previously identified to enhance circRNA synthesis using an circmCherry-expression screening system^15^. Using previously published quantitative mass spectrometry (MS) datasets for HSV-1 primary infection^53^ and KSHV reactivation^54^ we evaluated changes in protein abundance for our list of 27 predicted RBP interactors. Lytic infection drives host shut off resulting in a global decrease in host peptides^53,54^. Despite this, four (ACO1, HNRNPH2, SART3, SRSF7) and 13 RBPs (FUS, G3BP2, HNRNPC, HNRNPCL1, HNRNPH2, HNRNPK, KHDRBS1, MBNL1, PABPC1, PABPC4, SART3, SRSF7, ZC3H) were upregulated during HSV-1 infection and KSHV lytic reactivation, respectively. Whether these upregulated RBPs are responsible for coordinating viral circRNA synthesis remains to be determined. Ultimately our analysis identifies distinct cis-elements coordinating viral and host circRNA synthesis and begs the question—are viral circRNAs spliceosome products?

### Impact of spliceosome inhibition on viral circRNA synthesis

To perturb the spliceosome we treated HSV-1 (MRC-5 lytic infection) and KSHV (iSLK-BAC16 lytic reactivation) models with pladienolide B (PB) and isoginkgetin (IGG) which prevent formation of the major spliceosome A or B complex, respectively (Fig. 4A). We assessed cell viability and splicing inhibition in our models to determine the appropriate concentration of inhibitor to use to maximize inhibition and minimize off-target effects (Sup Fig. 4-1). Based on these assays we treated our models with 30 uM IGG and 25 nM PB and assessed viral transcription and replication. These drugs successfully inhibited forward splicing for viral transcripts as assessed by qPCR using exon-exon (mature mRNA) or exon-intron (pre-mRNA) specific primers and quantitation of RNA-Seq forward splice junctions (FSJ) (Fig. 4B). Spliceosome inhibition resulted in reduced HSV-1 and KSHV genome replication as well as HSV-1 infectious progeny (Fig. 4C). We expect this is due to decreased levels of the HSV-1 elongation factor (US1 or ICP22) and KSHV RNA binding protein (ORF57) (Fig. 4B) which are products of forward splicing.

**Fig. 4.**
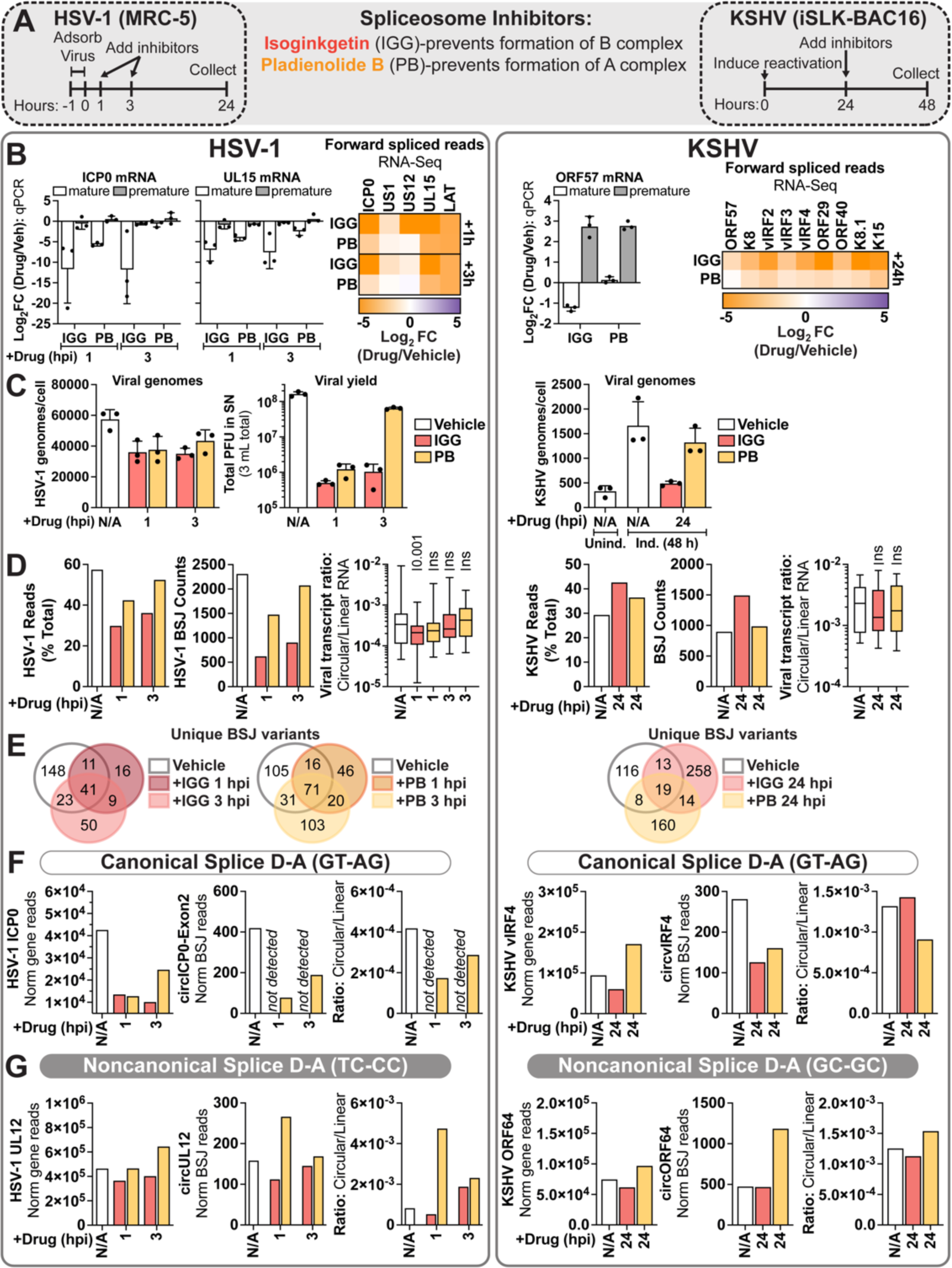
Impact of spliceosome inhibition on viral circRNA synthesis. Infection models were treated with spliceosome inhibitors, isoginkgetin (IGG) and pladienolide B (PB). HSV-1 infected fibroblasts were treated with inhibitors at 1 or 3 hours post infection and collected at 24 hours post infection. iSLK-BAC16 were treated with inhibitors at 24 hours after lytic reactivation and collected after 24 hours of inhibitor treatment. B, D-G) RNA-Seq was performed and normalized to ERCC spike-in reads, data is the average of biological duplicates. B) Forward spliced viral transcripts were assessed by qPCR (n=3) or RNA-Seq (n=2). Data is plotted as the log_2_ fold change (inhibitor/vehicle). C) qPCR assessment of genome quantity (n=3), plotted as the number of viral genomes per cell. HSV-1 infectious progeny (n=3) was measured by plaque assay and plotted as total plaque forming units (PFU) in the supernatant. Each point is a biological replicate, column bars are the average, and error bars are standard deviation. D) Sequencing overview for all viral mapped reads, as percent total reads (% Total). The number of total BSJ counts is reported for high-confidence viral circRNAs. Circ/Linear ratios for BSJ variants identified in all conditions are plotted as box and whisker plots. Wilcoxon paired t-tests were used to compare inhibitor treatment to vehicle. p-values are reported above. E) Overlapping identity of high-confidence viral circRNAs called in dataset. F-G) Impact of splicing inhibition for instances of viral circRNAs which use canonical or noncanonical splice donor-acceptors.

To determine a global picture of splicing changes we performed RNA-Seq and quantified viral circRNAs using CHARLIE (Fig. 4D-G). For HSV-1 there was a decrease in BSJ counts after inhibitor treatment, however this decrease was proportional to global changes in read composition (Fig 4D-E). For KSHV there was no decrease in the amount of BSJ reads or variants detected after spliceosome inhibition (Fig 4D-E). We calculated the ratio of circular/linear reads for a subset of circRNAs expressed in vehicle and inhibitor treated conditions. Using this metric, we can evaluate how perturbation alters the likelihood a given gene gives rise to a circRNA. Inhibitor treatment only caused a significant shift in circ/linear ratios for HSV-1 +IGG 1 hpi (Fig 4D). All other conditions did not result in a significant change in circRNA synthesis. In Fig 4F and G we highlight a subset of viral circRNAs for each model, one with and one without canonical splice DA elements. The circRNA species which arises from the middle exon of HSV-1 ICP0 is undetectable or decreased after spliceosome inhibition (Fig 4F). By comparison, the circRNA species colinear to UL12 is upregulated after spliceosome inhibition (Fig 4G). KSHV circvIRF4 had a decrease in BSJ reads after spliceosome depletion, however when plotted as circ/linear ratio there was minimal change in circRNA synthesis (Fig 4F). KSHV circORF64 BSJ levels were unchanged (IGG) or increased (PB) by spliceosome inhibition, with no change in circ/linear ratios when compared to vehicle (Fig 4G). To corroborate our KSHV findings we performed siRNA depletion of a core component of the spliceosome, *PRPF8* (Supplementary Fig 4-2A). After *PRPF8* depletion we observed a decrease in host and viral forward splicing (Supplementary Fig. 4-2B). By comparison, viral circRNAs (circvIRF4, circPAN, circK2) were unaffected by PRPF8 depletion (Supplementary Fig. 4-2C). In summary we find the majority of viral circRNAs to be unaffected by spliceosome inhibition. The notable exception would be HSV-1 circICP0-exon2, which was undetected after inhibitor treatment (Fig 4F). This data supports a model in which most viral back splicing is mediated by machinery other than the major spliceosome or a model in which viral back splicing is favored over forward splicing when spliceosome activity is limited.

### Impact of key lytic effectors on viral circRNA synthesis

While herpesviruses replicate in the nucleus and rely on the host machinery for transcription, splicing, and RNA export, they express viral proteins which target and modulate these pathways. To investigate how these unique effectors may influence circRNA biogenesis, we employed knockout viruses and inhibitors that target distinct phases of the lytic life cycle. These included viral transcription factors (HSV-1 ICP4 and ICP22, KSHV ORF24, MHV68 ORF50) and RNA binding proteins (HSV-1 ICP27, KSHV ORF57). We used cycloheximide (CHX) to block HSV-1 at the phase of immediate early transcription and DNA replication inhibitors (cidofivir or CDV, phosphonoacetic acid or PAA) to block viruses at the phase of immediate early and early transcription. We quantified viral genome replication and transcription, confirming that our infections echoed previously published phenotypes (Supplementary Fig. 5-1, Supplementary Fig. 5-2A-B)^55–62^. We performed RNA-Seq and quantified viral circRNAs using CHARLIE (Fig. 5, Supplementary Fig. 5-2C-D). Viral circRNA abundance and diversity echoed total read composition (Fig 5A, C), with the outlier being KSHV τιORF57. In the absence of the KSHV RBP, ORF57, there was a drastic decrease in the diversity of BSJ species (Fig 5C)–with a new, highly abundant circRNA appearing in the PAN locus (Fig 5D, F).

**Fig. 5.**
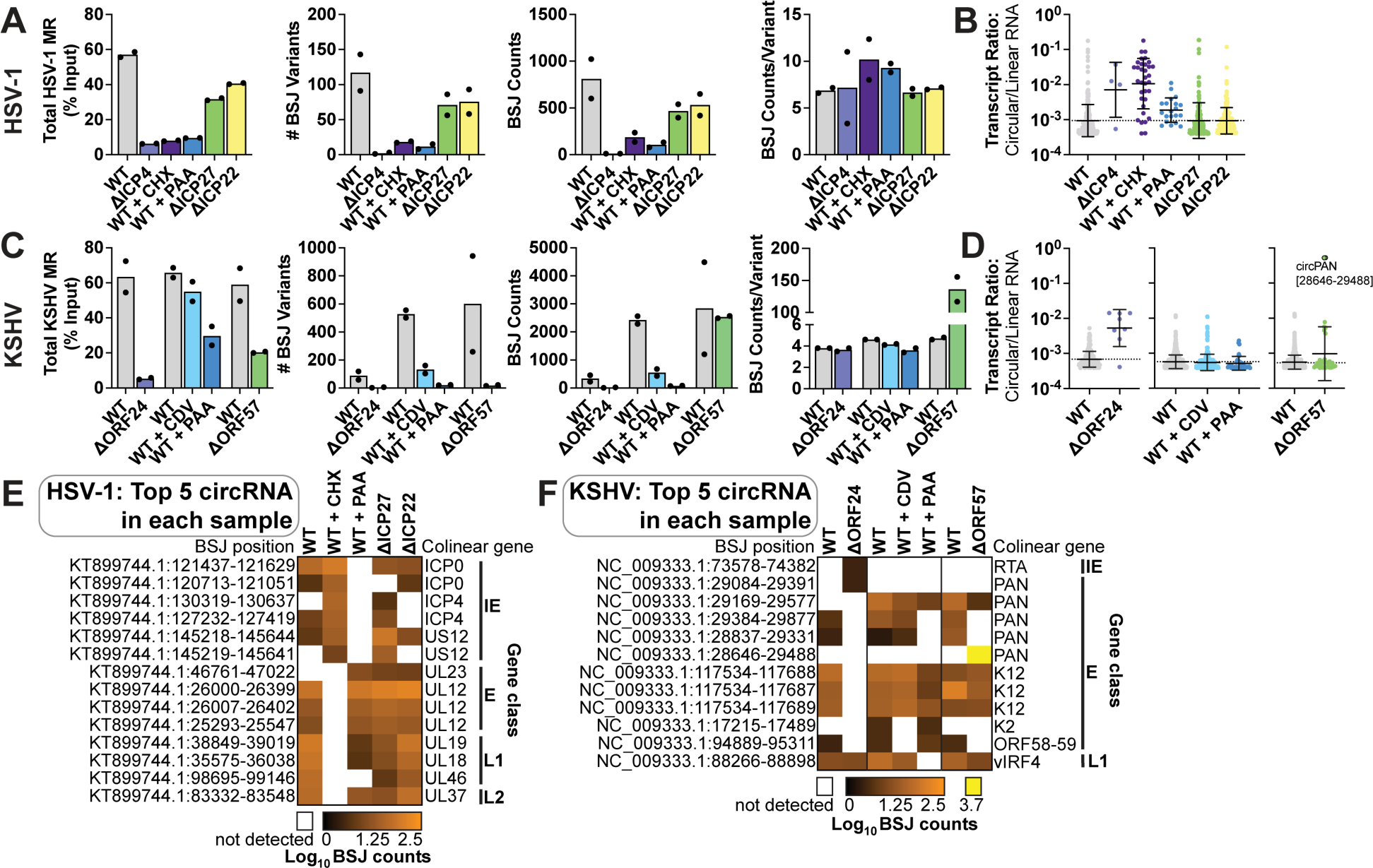
Impact of key lytic effectors on viral circRNA synthesis. RNA-Seq data from HSV-1 (MRC-5 cells infected at an MOI of 10 PFU/cell for 12 hours) and KSHV (iSLK-BAC16 induced with Dox and NaB for 3 days) infection. Experiments were performed with mutant cells and viruses which lack expression of the viral protein indicated. If indicated, cells were treated with cidofivir (CDV), phosphonoacetic acid (PAA), or cycloheximide (CHX) at 0 hours post infection (hpi). Data is the average of biological duplicates. Analysis is relative to a paired wildtype (WT) control. A, C) Sequencing overview for all viral mapped reads (MR), as percent total reads (% Total). The number of unique BSJ variants, total BSJ counts, or BSJ counts/variant is reported for high-confidence viral circRNAs. Each point is a biological replicate, column bars are the average. B, D) Circ/Linear ratios for high-confidence viral BSJ variants. Each dot represents a unique BSJ, cross-bars are the geometric mean and error bars are the geometric standard deviation. E) The top five most highly expressed circRNAs in each sample were merged to form heatmaps which report Log_10_ BSJ counts for biological duplicates. BSJ positions are reported on the left. Colinear genes and their gene class are reported on the right, with IE (immediate early), E (early), L1 (leaky late), L2 (true late).

To account for global transcriptional changes in these infections, we quantified the circ/linear transcript ratio for only actively transcribed regions of the genome. We observed a higher circ/linear ratio for viral genes in the absence of the viral transcription factors HSV-1 ICP4 and KSHV ORF24 (Fig 5B, D). As the number of circRNA species generated in these conditions is much lower than wildtype, we are uncertain of the significance of this shift. No MHV68 circRNAs were identified in the absence of the viral transcription factor, ORF50, thus circ/linear ratio could not be determined (Supplementary Fig. 5-2C-D). Inhibition of viral DNA replication had minimal impact on circ/linear ratios for HSV-1, KSHV, and MHV68 (Fig. 5A-D, Supplementary Fig. 5-2C-D). We observed no significant change in HSV-1 circRNA synthesis in the absence of the viral RBP (ICP27) or transcription elongation factor (ICP22) (Fig. 5E, Supplementary Fig. 5-3A). We plotted the top five most highly expressed BSJ variants from each condition and compared their levels (Fig. 5E-F). For HSV-1, we noted that the colinear genes which give rise to highly expressed circRNAs echoed the kinetic class licensed in each condition (Supplementary Fig. 5-3A). The shift in profile during HSV-1 infection leads us to suspect no specific viral effector tested was responsible for promoting circRNA biogenesis, and instead that any highly expressed viral gene has the potential to generate a circular isoform. For KSHV, we found only the viral RNA binding protein, ORF57, significantly altered viral circRNA profile.

### ORF57 regulation of circRNA synthesis

Next, we investigated the potential role that KSHV ORF57 may have in controlling circRNA biogenesis. As ORF57 is a viral RNA binding protein^63,64^ we first assessed if it directly bound circRNAs. We performed eCLIP (enhanced version of the crosslinking and immunoprecipitation) on iSLK-BAC16 reactivated for 24 hours by treating with Dox/NaB (Fig. 6A-B). ORF57 immunoprecipitated (IP) or size-matched input (input) samples were sequenced to generate 50 bp single-end reads. Forward and back splice junctions were quantified using STAR or CHARLIE. We observed a general enrichment for viral reads within ORF57 IP samples (Fig. 6A). ORF57 IP resulted in an enrichment for KSHV and host back spliced transcripts as well as host forward spliced transcripts (Fig. 6B). Of the 71 viral BSJ enriched after ORF57 IP, 59 were from species colinear with PAN (Supplementary Fig. 6-1A). Host genes with ≥2 BSJ variants enriched in the ORF57 IP were GLS, RMRP, MT-RNR2 (Supplementary Fig. 6-1B). These data demonstrate that ORF57 binds a subset of host and viral circRNAs within close proximity (50 nt) of the back spliced junction.

**Fig. 6.**
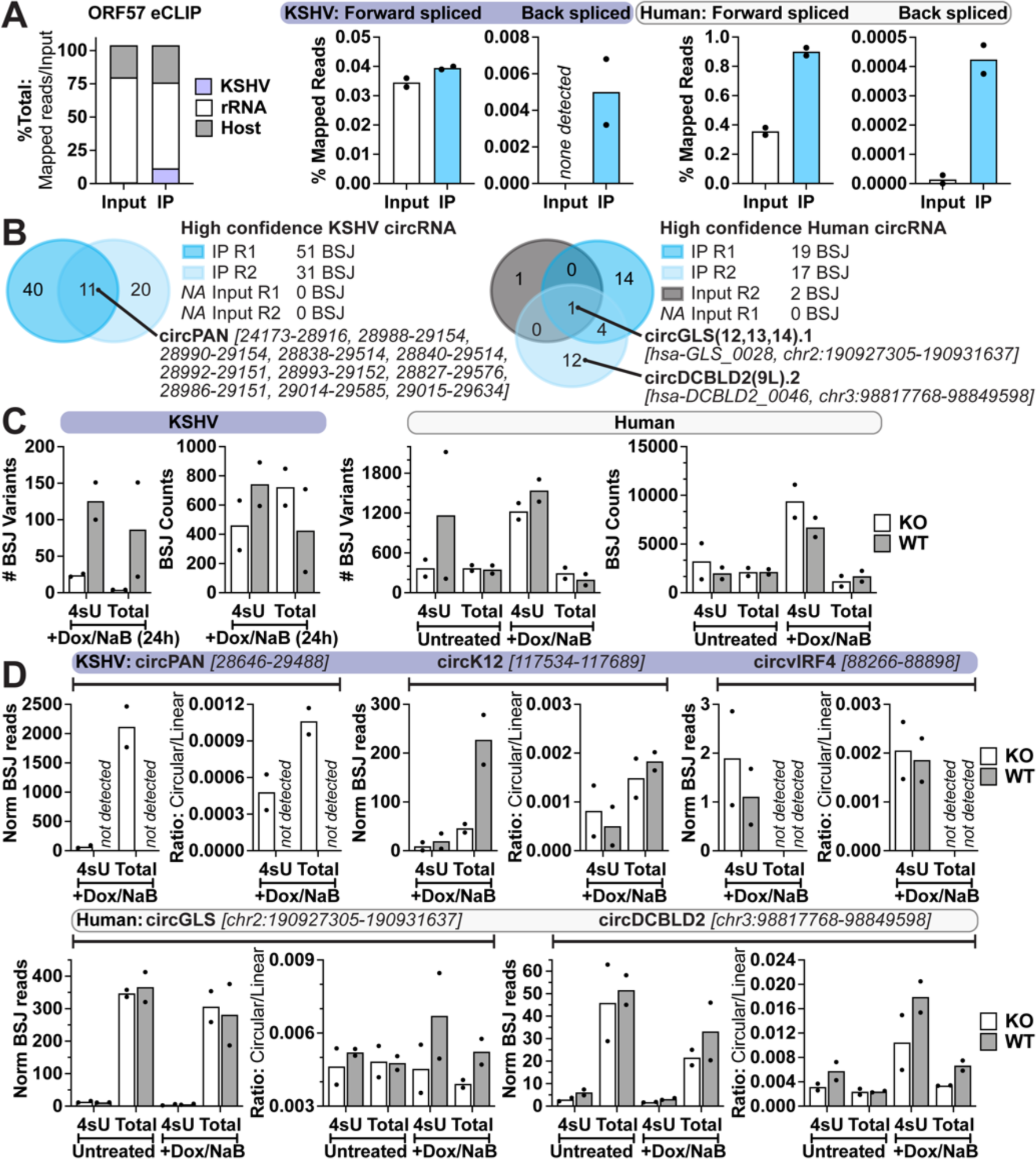
ORF57 regulation of circRNA synthesis. A-B) ORF57 eCLIP (n=2) was performed on iSLK-BAC16 reactivated for 24 h. “Input” is a paired, size-selected RNA-Seq wherein ORF57 immunoprecipitation (IP) was not performed. A) Reads mapped to ribosomal RNA (rRNA), human genes (excluding rRNA), or KSHV genes are plotted as a function of total reads. Uniquely mapped reads containing forward or back splice junctions were plotted relative to all reads mapped to their respective genome. Column bars are the average and data points are individual biological replicates. B) Overlapping identity of high-confidence circRNAs called in dataset. C-D) Bulk RNA-Seq (Total) and 4sU RNA-Seq (4sU) was performed on ϕλORF57 (KO) or iSLK-BAC16 (WT) cells with (24 h) or without (Unind.) reactivation. 4sU was added to media 15 minutes prior to the collection time point. BSJ counts are the average of biological duplicates and normalized as 4sU-Seq: Reads Per Kilobase per Million mapped reads (RPKM) or Total: relative to ERCC spike-ins. Each point is a biological replicate, column bars are the average. C) The number of unique BSJ variants and total BSJ counts is reported for all high confidence circRNAs. D) Normalized BSJ counts and circ/linear ratios for select circRNAs.

ORF57 possesses homologs in all human herpesviruses, many of which have been demonstrated to enhance co- and post-transcriptional gene regulatory activity^65,66^. We employed Total and Nascent RNA (4sU)-Seq to determine if ORF57 promotes circRNA accumulation by influencing circRNA synthesis or reducing transcript decay (Fig. 6C-D, Supplementary Fig. 6-2). We first evaluated our approach by analyzing linear viral transcripts (Supplementary Fig. 6-2). To compare transcriptional changes we plotted the log_2_ fold change (ΔORF57/WT) for total (x-axis) or nascent (y-axis) RNA levels. Linear KSHV transcripts at 24 hpi stratified as ORF57 enhanced post-transcriptionally (Supplementary Fig. 6-2B). PAN, ORF6-9, and ORF58-59 were the most down-regulated in the absence of ORF57. The notable exceptions to this were K15 and ORF75, which clustered as ORF57 repressed post-transcriptionally. These findings are consistent with a recent study demonstrating that ORF57 prevents premature accumulation of viral late genes by modulating the host RNA decay machinery^67^. This provides increased confidence that our method can successfully delineate ORF57 mediated co- or post-transcriptional regulatory effects.

We analyzed our data using CHARLIE to compare changes in host and circRNA levels between KO (1′ORF57) and WT (Fig 6C-D). We limited our analysis to uninduced and 24 hpi, to remove any confounding effects from DNA replication defects in KO cells (Supplementary Fig. 5-1A). We observed a greater diversity of viral BSJ variants in both 4sU and total RNA-Seq during WT infection (Fig. 6C). In the absence of ORF57, a novel, highly abundant (>500 raw BSJ reads) circPAN [28646:29488] was expressed; other circPAN species expressed during WT infection were largely absent (Supplementary Fig 6-3). This novel circPAN variant was at the 5’ end of PAN and did not contain the element for nuclear expression (ENE) (Supplementary Fig 6-3A). circPAN [28646:29488] expression was validated by qPCR using divergent primers and a probe that spans the exact BSJ sequence identified in our RNA-Seq (Supplementary Fig 6-3B). We examined expression of other KSHV circRNAs, circK12 and circvIRF4, and saw little difference in expression when comparing KO and WT. We support a model in which ORF57 regulates circularization of the PAN transcript, favoring a cluster of many diverse species and preventing synthesis of circPAN [28646:29488]. We noted an increase in host circRNAs (BSJ counts) at 24 hpi in our 4sU-Seq data (Fig 6C). This trend suggests that lytic infection induces synthesis of host circRNAs. Thus, we evaluated expression changes for host circRNA enriched in our ORF57 eCLIP IP. We found circGLS and circDCBLD2 to be upregulated at 24 hpi, and this upregulation was abrogated in KO cells (Fig 6D). Host circRNA increase was more pronounced in 4sU-Seq data, suggesting ORF57 promotes synthesis rather than preventing decay. In summary, KSHV ORF57 promotes circRNA synthesis for a subset of viral and host circRNAs during lytic infection.

## DISCUSSION

Herein, we performed comparative circRNA profiling for HSV-1, KSHV, and MHV68 during primary lytic infection, latency, and reactivation. CircRNAs are a novel class of transcripts, with long half-lives and low immunogenicity^28^. As alternative splicing products, circRNAs extend transcriptional density without expanding genome space. These qualities certainly seem advantageous to a pathogen in the context of viral infection. We identified thousands of viral circRNAs tiling the viral genome. We generated an annotated resource table with the identity, position, and sequence of all circRNAs detected in the array of models tested (Supplementary Dataset 1). Select viral circRNAs were highly abundant (HSV-1 circUL44, KSHV circPAN) reaching levels within 100-fold of an abundant host housekeeping gene, *GAPDH*. Recent work has shown that viral circRNAs can serve as translational templates^22^, be packaged in virions and extracellular vesicles^23^, and modulate host cell division and survival^17,29^. While abundance does not always equate biological relevance, the high-copy number of these viral circRNAs certainly begs the question of molecular function.

Our profiling identified circRNA species expressed during all phases of the viral life cycle, namely HSV-1 circLAT, KSHV circK12, and MHV68 circORF66. We performed *in silico* prediction of circRNA-miRNA-mRNA networks for these circRNAs (Supplementary Fig. 1-6). CircRNAs generally function to regulate gene expression and one mechanism by which this occurs, is miRNA sponging^28,48^. In this way, circRNAs repress miRNA-mediated mRNA degradation. Overrepresentation analysis on downstream mRNA targets in the network found that circLAT may modulate cellular senescence by targeting hsa-miR-124-3p. The parental gene, LAT, has previously been shown to influence neuronal senescence and apoptosis, a finding that may establish a link between HSV-1 infection and neurodegenerative diseases^68,69^. KSHV circK12 was predicted to influence mTOR signaling with *AKT3* as a potential downstream mRNA target (Supplementary Fig. 1-6). A number of KSHV viral proteins have established roles in modulating the PI3K/AKT/mTOR axis^70^, as the pathway controls B lymphocyte proliferation and development. One can envision how a non-immunogenic, long-lived, circRNA species with the potential to be expressed during all phases of the life cycle would be an advantageous molecule to also target this axis. These *in silico* predictions provide testable hypotheses for functional studies of circLAT and circK12 during infection.

CircRNA biogenesis has been largely characterized for model organisms, such as human, mouse, and drosophila. In these models, back splicing is catalyzed by the spliceosome and promoted by RNA binding proteins (RBPs) and tandem repeats which mediate interaction of flanking sequences^11–15^. As an alternative splicing event, circRNAs rely on canonical splice DA signals, such as GT-AG^11,12^. A spliceosome-independent mechanism for circRNA synthesis was recently identified in archaea (*H. volcanii*) and metazoa (*D. melanogaster*)^71,72^. Here, the tRNA splicing machinery creates circularized tRNA introns (tricRNA) via tRNA splicing endonuclease (TSEN)-mediated intron cleavage and RTCB catalyzed 3’ to 5’ RNA ligation. tRNA introns are not flanked by splice DA signals, instead they are recognized by a secondary structure element, called a bulge-helix-bulge motif^73^. Additional methods for spliceosome-independent circularization have been published for *in vitro* circRNA synthesis^74,75^. These leverage either chemical or enzymatic ligation. In the case of enzymatic ligation, T4 DNA ligase (with a complementary ssDNA splint), T4 RNA ligase, or RTCB can circularize RNA.

Spliceosome dependent back splicing was assumed to also apply to virus-derived circRNAs, including KSHV circ-vIRF4 and EBV circRPMS1. These viral circRNAs are colinear with multi-exon viral genes and coordinated by flanking splice DA sites^18,76^. However, the structure of these viral genes is not the norm, but rather the exception. Most viral transcripts mimic premature-tRNA in that they are short (∼500 nt), usually single exon (lack splice donor-acceptor sites), and highly structured (high GC-content or repeat elements). Herein, we report that the majority (>90%) of viral circRNAs lack canonical splice donor-acceptor sites (Fig. 3) and do not reside in multi-exon genes (Fig. 2). Most viral circRNAs were unaffected by spliceosome inhibition or depletion (Fig 4, Supplementary Fig. 4-2). HSV-1 circRNAs tile the viral genome and appear to be coordinated by high GC content in flanking regions (Fig 2B, Fig 3C). This data supports a model in which viral back splicing is mediated by machinery other than the major spliceosome, or a model in which viral back splicing is favored over forward splicing when spliceosome activity is limited. The latter model has previously been proposed for circRNAs expressed from the host genome^77^. As viral back splicing occurs independent of the canonical splice donor-acceptor (GT-AG), we support the first hypothesis for viral circRNA biogenesis. This is of special interest for HSV-1 which only expresses five spliced mRNAs and inhibits the spliceosome^50,51^. The potential for RNA ligase-dependent circRNA synthesis in humans has yet to be explored; although the tRNA splicing machinery is highly conserved, and humans express a homolog of RTCB and the TSEN complex. Additionally, a human 5’ to 3’ RNA ligase (C12orf29) was recently discovered^78^, with potential to generate circular products. We posit that these or other RNA ligases may be capable of generating circular viral RNA products although further work is necessary to determine the veracity of this model.

CircRNA maturation is frequently mediated by host RNA binding proteins, which bind adjacent to the 5’ and 3’ back splice junction and bridge the gap. In line with this, we tested the potential contribution of the KSHV encoded RBP (ORF57) in promoting viral back splicing. We used complimentary techniques (eCLIP, 4sU- and Total-RNA-Seq) and determined that ORF57 directly bound near back splice junctions in circPAN and shifted the circRNA species expressed from this locus (Fig. 6). Additionally, we found instances of host circRNAs bound by ORF57 (circGLS, circDCBLD2) with increased nascent levels only in the presence of ORF57. By comparing 4sU and Total RNA-Seq, we concluded that ORF57’s function is independent from its traditionally published role in modulating RNA decay. Whether ORF57 directly impacts circRNA synthesis through changing the rate of parental gene expression or directly impacting back splicing rates is unclear. A recent publication demonstrated that ORF57 enhanced accumulation of the host circRNA, circHIPK3^79^. Our study compliments this, extending the observation to additional viral and host circRNAs.

Herein we find that lytic infection results in synthesis of abundant and diverse viral circRNA species; with a subset expressed in latency and during lytic reactivation. Comparative profiling of host and viral circRNAs after perturbation of key factors— including the spliceosome, viral transcriptional activators, and RNA binding proteins— identified divergent phenotypes and suggests there are distinct mechanisms of synthesis. As viral circRNAs have only been studied in the last five years, our work has the potential to identify new factors which contribute to herpesvirus replication, persistence, and tumorigenesis. This in turn may aid in identification of novel clinical biomarkers, drug targets, and development of antiviral compounds.

## METHODS

### Cells and Viruses

Vero (ATCC #CCL-81) and Vero-based HSV-1 complementing cell lines (E5, E11) were maintained in DMEM (Gibco #11965-092) supplemented with 5% fetal bovine serum (FBS), 1 mM sodium pyruvate, 2 mM l-glutamine, 100 units/mL penicillin-streptomycin. MRC-5 (ATCC #CCL-171) were maintained in DMEM (Gibco #11965-092) supplemented with 10% FBS, 1 mM sodium pyruvate, 2 mM l-glutamine, 100 units/mL penicillin-streptomycin. NIH 3T3 (ATCC #CRL-1658) and NIH 3T12 (ATCC #CCL-164) were maintained in DMEM (Corning # 10-017-CV) supplemented with 8% FBS, 2 mM l-glutamine, 100 units/mL penicillin-streptomycin. A20 HE-RIT B cells harbor a recombinant MHV68 expressing a Hygromycin-eGFP cassette^80^ and these cells express MHV68 RTA under the control of a doxycycline-inducible promoter^81^. A20 HE-RIT were maintained in RPMI supplemented with 10% FBS, 100 units/mL penicillin-streptomycin, 2 mM l-glutamine, 50 uM beta-mercaptoethanol, 300 ug/ml hygromycin B, 300 ug/mL gentamicin, and 2 ug/mL puromycin. iSLK-BAC16 renal carcinoma cells harbor a recombinant KSHV expressing a Hygromycin-eGFP cassette and these cells express KSHV RTA under the control of a doxycycline-inducible promoter^82^. iSLK-BAC16 (WT, ΔORF24, ΔORF57) were maintained in DMEM (Gibco #11965-092) supplemented with 10% Tet-Approved FBS, 50 ug/mL hygromycin B, 0.1 mg/mL gentamicin, 1 ug/mL puromycin, 100 units/mL penicillin-streptomycin. HDLEC (PromoCell #C-12216) and HUVEC (Lonza #C2519A) were maintained in EBMTM-2 Basal Medium (Lonza #CC-3156) supplemented with EGMTM-2 SingleQuots Supplements (Lonza #CC-4176). We thank Neal DeLuca (n12, 5dl1.2, n199, E5, E11) and Britt Glaunsinger (iSLK-BAC16, iSLK-BAC16 ΔORF24) for their kind gift of viruses and cells. Cells and viruses used in this study are listed in Supplementary Table 3.

### Oligos

A list of all oligos used is located in Supplementary Table 4.

### Virus Stock Preparation and Titration

**HSV-1.** Vero-based cell lines were infected at a low MOI (∼0.01 PFU/cell) and harvested when cells were sloughing from the sides of the vessel. Supernatant and cell fractions were collected and centrifuged at 4,000 x g 4°C for 10 minutes. The subsequent supernatant fraction was reserved. The pellet fraction was freeze (−80°C 20 min)/thawed (37°C 5 min) for three cycles, sonicated for 1 minute (70% power, water bath), and centrifuged at 2,000 x g 4°C for 10 minutes. The final virus stock was composed of the cell-associated virus and reserved supernatant virus fractions. KOS, strain 17, and n199 were prepared and titered in Vero cells. Other virus stocks were prepared and titered in the following Vero-based complementing cell lines: E5 (ICP4+, n12-complementing) and E11 (ICP4/ICP27+, 5dl1.2-complementing).

**KSHV.** iSLK-BAC16 cells were induced with 1 ug/mL doxycycline and 1 mM sodium butyrate for 3 days. Cell debris was removed from the supernatant fraction by centrifuging at 2000 x g 4°C for 10 minutes and filtering with a 0.45 PES membrane. Virus was concentrated after a 16,000 x g 4°C 24 hour spin and resuspended in a low volume of EGM2 media (approx. 1000-fold concentration). To assess viral infectivity, VEC or LEC cells were infected with serial dilutions of the BAC16 stock and assessed using Flow Cytometry for GFP+ cells at 3 days post infection. BAC16 contains a constitutively expressed GFP gene within the viral genome. Based on these assays, BAC16 stock was used at a 1:60 dilution, resulting in 35% infection for HUVEC (MOI 0.25) and 70% infection for LEC (MOI 1).

**MHV68.** NIH3T12 based cell lines were infected at a low MOI with H2B-YFP until 50% cytopathic effect was observed. Infected cells and conditioned media were dounce homogenized and clarified at 600 x g 4°C for 10 minutes. Clarified supernatant was further centrifuged at 3,000 x g 4°C for 15 min and then 10,000 x g 4°C for 2 hrs to concentrate ∼40-fold in DMEM. H2B-YFP was prepared and titered using plaque assays in NIH 3T12 cells. The recombinant MHV68 ORF50.stop virus was produced and titered using plaque assays in CS-RTA4 3T12 cells^83^.

### *De Novo* Infection

**HSV-1.** Confluent MRC-5 cells were infected with 10 plaque forming units (PFU) per cell. Virus was adsorbed in PBS for 1 hr at room temperature. Viral inoculum was removed, and cells were washed quickly with PBS before adding on DMEM media supplemented with 2% FBS. 0 hour time point was considered after adsorption of infected monolayers when cells were place at 37°C to incubate. If indicated, cells were treated with 75 ug/mL Cycloheximide (Sigma #C4859) or 300 ug/mL Phosphonacetic Acid (Sigma #284270) at 0 hr post infection.

**KSHV.** VEC or LEC cells were infected with BAC16 at an approximate MOI of 0.25 (30% cells infected) or 1 (70% cells infected), respectively. Virus was adsorbed in a low volume of media for 8 hr at 37°C, after which viral inoculum was removed and replaced with fresh media. 0 hour time point was when virus was added and cells were first placed at 37°C to incubate.

**MHV68.** Subconfluent NIH3T12 fibroblasts were infected with 5 PFU per cell. Virus was adsorbed in a low volume of DMEM media supplemented with 8% FBS for 1 hr at 37°C, prior to overlay with fresh media. 0 hour time point was when virus was first added and cells were placed at 37°C to incubate. If indicated, cells were treated with 100 ug/mL Phosphonacetic Acid at 1.5 hr post infection.

### HSV-1 Mouse Infections

Female 8-week-old BALB/cAnNTac mice were infected with 10^5^ PFU HSV-1 (strain 17) via the ocular route. After 4-5 weeks, latently infected trigeminal ganglia were harvested and immediately processed or explanted into culture to induce viral reactivation. Trigeminal ganglia explants were cultured (DMEM/1% FBS) for 12 hours at 37°C/5% CO_2_ in the presence of vehicle (DMSO), 100 uM acyclovir, or 2 uM JQ1 (Cayman Chemical CAS: 1268524-70-4). Pools of 6 ganglia were homogenized in 1 ml TriPure isolation reagent (Roche) using lysing matrix D on a FastPrep24 instrument (3 cycles of 40 seconds at 6 m/s). 0.2 ml chloroform was added for phase separation using phase lock gel heavy tubes and RNA isolation from the aqueous phase was obtained by using ISOLATE II RNA Mini Kit (Bioline). RNA quality was verified with Agilent 2100 Bioanalyzer System using RNA Nano Chips (Agilent Technologies). All animal care and handling were done in accordance with the U.S. National Institutes of Health Animal Care and Use Guidelines and as approved by the National Institute of Allergy and Infectious Diseases Animal Care and Use Committee (Protocol LVD40E, T.M.K.).

### MHV68 Mouse Infections

Mixed bone marrow chimeric mice generated by reconstitution of CD45.1 C57/BL6 recipient mice with bone marrow from CD45.2 *CD19^cre/+^*STAT3 wild-type mice and *CD19^cre/+^Stat3^f/f^tdTomato^stopf/f^*B-cell STAT3 knock-out mice were infected by intraperitoneal injection with 1,000 PFU MHV68 H2B-YFP in 0.5 ml under isoflurane anesthesia. At 16 days post-infection (dpi), mouse spleens were homogenized, treated to remove red blood cells, and enriched for B-cells (Pan B-cell isolation kit; STEMCELL, Vancouver, BC, Canada). YFP+ infected and YFP-uninfected GL7+CD95+ germinal center (GC) B-cells were sorted and collected from tdTomato- (*CD19^cre/+^* wild-type) CD3-B220+CD45.2+ B-cells. 5 x 10^4^ cells sorted from pooled mouse splenocytes were spun down, resuspended in 50 µl of TRIzol, and stored at −80°C. GENEWIZ (South Plainfield, NJ) performed RNA extraction, quality control, library preparation, and Illumina sequencing. Additional experimental details are included in Hogan *et al.* (2023)^84^. All mouse experiments were performed in accordance with protocols approved by the National Cancer Institute Animal Care and Use Committee.

### Lytic Reactivation

**KSHV.** Subconfluent monolayers of iSLK-BAC16 (WT, ΔORF24, ΔORF57) were induced with 1 ug/mL doxycycline, 1 mM sodium butyrate in DMEM media supplemented with 2% Tet-approved FBS. 0 hour time point was when induction media was added and cells were first placed at 37°C to incubate. If indicated, cells were treated with 100 uM Cidofovir (Sigma) or 70 ug/mL Phosphonacetic Acid (Sigma) at 0 hr post induction.

**MHV68.** One day prior to induction, A20 HE-RIT cells were subcultured at a 1:3 dilution in media lacking antibiotics. Cells were seeded subconfluently and induced for 24 hours with RPMI media containing 5 ug/ml Dox and 20 ng/ml TPA.

### RNase R Confirmation of Viral CircRNAs

**HSV-1.** Total RNA was isolated from cells using the Direct-zol RNA MiniPrep Kit (Zymo), following manufacturer’s instructions. 7 ug total RNA was combined with 1x E-PAP Buffer, 2.5 mM MnCl2, 1 mM ATP, 40 Units RNase Inhibitor (ThermoFisher), and 8 Units E-PAP (PolyA Tailing Kit, ThermoFisher). Poly-A tailing reactions were incubated at 37°C for 10 minutes. RNA was acid-phenol chloroform extracted, ethanol precipitated, and resuspended in nuclease-free water. PolyA-Tailed material was combined with 20 Units RNase Inhibitor (ThermoFisher), 20 Units RNase R (Lucigen), 100 mM LiCl, 20 mM Tris-HCl pH 8.0, 0.1 mM MgCl_2_. RNA was RNase R digested at 37°C for 30 minutes, following clean-up with the RNA Clean and Concentration Kit (Zymo). RNA was reverse transcribed with random decamers using ReverTra Ace-α qPCR RT Master Mix (Toyobo). cDNA was measured using quantitative PCR (qPCR) with either divergent (circRNA) or convergent (mRNA) primers.

**KSHV and MHV68.** Total RNA was isolated from cells using the Direct-zol RNA MiniPrep Kit (Zymo), following manufacturer’s instructions. 3 ug total RNA was combined with 20 Units RNase Inhibitor (ThermoFisher), 20 Units RNase R (Lucigen), 100 mM KCl, 20 mM Tris-HCl pH 8.0, 0.1 mM MgCl_2_. RNA was RNase R digested at 37°C for 30 minutes, following clean-up with the RNA Clean and Concentration Kit (Zymo). RNA was reverse transcribed with random decamers using ReverTra Ace-α qPCR RT Master Mix (Toyobo). cDNA was measured using quantitative PCR (qPCR) with either divergent (circRNA) or convergent (mRNA) primers.

### Splicing Inhibitor Assays

At various times relative to HSV-1 infection (MRC-5 infected with KOS at MOI of 10) or KSHV reactivation (iSLK-BAC16 + 1mM NaB 1 ug/mL Dox), supernatant was removed and replaced with media containing 30 uM Isoginkgetin (CAS #548-19-6, Sigma #416154) or 25 nM Pladienolide B (CAS #445493-23-2, SantaCruz #sc-391691). When indicated (vehicle), DMSO (0.1%) was added to media. Total RNA was collected at either 24 hours post infection (HSV-1, inhibitors added +1 or +3 hpi) or 48 hours post induction (KSHV, inhibitors added 24 hpi). Total RNA was isolated from cells using the Direct-zol RNA MiniPrep Kit (Zymo #R2053), following manufacturer’s instructions. For detection of unspliced targets—18S rRNA, pre-ORF57, pre-ICP0, pre-UL15—RNA was reverse transcribed with random decamers using the ReverTra Ace-α qPCR RT Master Mix (Toyobo #FSQ-201). For detection of spliced targets—mature-ORF57, mature-ICP0, mature-UL15—RNA was reverse transcribed with polydT oligos (Invitrogen #N8080128) using the High-Capacity cDNA Reverse Transcription Kit (Applied Biosystems #4374966). cDNA was measured using qPCR and primers which bound within intronic regions (unspliced) or spanned exon to exon boundaries (spliced). To measure HSV-1 infectious viral progeny, the following was performed. At 1 or 3 hours post infection, supernatant was removed and replaced with media containing 30 uM IGG or 25 nM PB. At 24 hours post infection, supernatant was collected and freeze-thawed 3 times. Vero cells were seeded at a density of 10^6^ cells/well in a 6-well dish and infected with serial diluted supernatant. Viral yield was determined by plaque assay and plotted as total PFU for the entire volume of supernatant collected.

### Measuring viral genomes

The cell fraction was isolated from infection models. Cell pellets were washed with 1x PBS and lysed using 0.5% SDS, 400µg/mL proteinase K, 100 mM NaCl. Samples were incubated at 37°C for 12-18 hours and heat inactivated for 30 minutes at 65°C. DNA samples were serial diluted 1:1000 and measured using qPCR with primers specific to HSV-1 UL23, KSHV ORF6, MHV68 ORF50, human GAPDH, or mouse GAPDH. Standard curves were generated using purified genomic stocks (HSV-1 BAC, KSHV BAC, human genome Promega #G1471, mouse genome Promega #G3091), or purified plasmids (MHV68 ORF50). Absolute copy number of genomic stocks was determined using ddPCR. Values were plotted as follows: 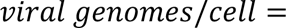,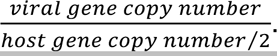

### Ribominus Total RNA-Seq

Total RNA was isolated from cells using the Direct-zol RNA MiniPrep Kit (Zymo), following manufacturer’s instructions. ERCC spike-in controls (ThermoFisher) were added to 500-1000 ng of total RNA. RNA was sent to the NCI CCR-Illumina Sequencing facility for library preparation and sequencing. RNA was ribominus selected and directional cDNA libraries were generated using either Stranded Total RNA Prep with Ribo-Zero Plus (Illumina # 20040525) or TruSeq Stranded Total RNA Ribo-Zero Gold (Illumina #RS-122-2303). 2-4 biological replicates were sequenced for all samples. Sequencing was performed at the NCI CCR Frederick Sequencing Facility using the Illumina NextSeq 550 or Illumina NovaSeq SP platform to generate 150 bp PE reads.

### 4sU-Sequencing

iSLK-BAC16 cells were uninduced or induced with 1 ug/mL Doxycycline 1 mM Sodium Butyrate. At indicated times relative to induction, 10 mM 4sU (Sigma) was added to cell culture medium. 15 minutes post-4sU addition, cells were collected and RNA extracted using Directzol RNA MiniPrep Plus (Zymo). 40-50 ug total RNA was biotinylated in 10 mM Tris pH 7.4, 1 mM EDTA, 0.2 mg/mL EZ-link Biotin-HPDP (ThermoFisher). Unbound biotin was removed by performing a chloroform:isoamyl alcohol extraction using MaXtract High Density tubes (Qiagen). RNA was isopropanol precipitated and resuspended in water. Biotinylated RNA was bound 1:1 to Dynabeads My One Streptavidin T1 equilibrated in 10 mM Tris pH 7.5, 1 mM EDTA, 2 M NaCl. Bound beads were washed three times with 5 mM Tris pH 7.5, 1 mM EDTA, 1 M NaCl. 4sU-RNA was eluted with 100 mM DTT and isolated using the RNeasy MinElute Cleanup Kit (Qiagen). RNA was sent to the NCI CCR-Illumina Sequencing facility for library preparation and sequencing. Pulldown RNA was ribominus selected using the NEBNext rRNA Depletion Kit v2 (NEB # E7400L) and RNA-Seq libraries were generated using the NEBNext® Ultra II Directional RNA Library Prep Kit (NEB # E7760L). Two biological replicates were sequenced using the Illumina NextSeq 550 platform to generate 150 bp PE reads.

### eCLIP Sequencing

Subconfluent monolayers of iSLK-BAC16 were induced for 24 hours with 1 ug/mL Doxycycline, 1 mM Sodium Butyrate in DMEM media supplemented with Tet-approved FBS. ORF57 eCLIP was performed by Eclipse Bioinnovations (San Diego, CA) following the published protocol^85^. 10% of ORF57 immunoprecipitations, and 1% of inputs were run on NuPAGE 4-12% Bis-Tris protein gels and transferred to a nitrocellulose membrane. Membranes were probed using ORF57 (1:6,000 dilution) primary antibody and EasyBlot anti Rabbit IgG (1:10,000 dilution) and imaged with C300 Imager using Azure Radiance ECL. ORF57 was detected by western blot running around 75 kDa, so IP and size-matched input were taken from 51 kDa to 126 kDa regions. For these assays, two independent biological replicates were performed using an affinity purified anti-ORF57 polyclonal antibody^64,86^. An IgG control antibody was tested in parallel, but the yield from this was low, so all subsequent comparisons were performed relative to the size-matched input control.

## BIOINFORMATIC ANALYSIS

### *De novo* circRNA annotation

Illumina SRS data was analyzed using Circrnas in Host And viRuses anaLysis pIpEline (CHARLIE) v.0.9.0, source code available at https://github.com/CCBR/CHARLIE. The workflow is briefly as follows. RNA-Sequencing reads were trimmed using CutAdapt^87^. Trimmed reads were mapped using STAR (2-pass mapping) to concatenated genome assemblies which contain the host (hg38 or mm39) + virus (KT899744.1, NC_009333.1, MH636806.1) + ERCC spike-in controls^44^. Our pipeline combines five previously published tools for circRNA discovery, CIRCexplorer2^39^, CIRI2^40^, DCC^41^, circRNAFinder^42^, and find_circ^43^. BSJ calls must meet the following criteria: ≥3 counts, 18 nucleotides (nt) mapped on both sides of the BSJ, circRNA splice donor acceptor distance ≥150 nt, viral circRNA splice donor acceptor distance ≤5000 nt. Viral circRNA splice donor-acceptor distance was limited to 5 kb to prevent artefactual calls caused by the isomeric structure of the HSV-1 genome. For PE data, BSJ were further filtered, requiring the mate be contained within the 5’ and 3’ junction of the chimeric mapped read. High confidence circRNAs were those in agreement between CIRCexplorer2 STAR^44^ and BWA^45^ mapping outputs and called in at least one other tool. circRNAs are labeled based on their colinear genes, chromosome, and 5’ and 3’ BSJ positions. Genomic BSJ positions were used to determine circRNA species overlapping between conditions. To visualize data, STAR mapped BAMs were filtered for “linear” and “circular” reads. “Linear” BAMs included all reads, except those spanning BSJ. “circular” BAMs included reads spanning a BSJ. Bigwigs were generated using deepTools2 bamcoverage and bigwigCompare with 1 bp bins^88^. Transcript traces were visualized using Integrative Genomics Viewer (IGV)^89^.

### Transcriptome analysis

#### Standard gene expression analysis

RNA-Sequencing reads were trimmed using Cutadapt^87^ and the following parameters: --pair-filter=any, --nextseq-trim=2, --trim-n, -n 5, --max-n 0.5, -0 5, -q 20, -m 15. Trimmed reads were mapped using STAR^44^ with 2-pass mapping to concatenated genome assemblies which contain the host genome (hg38 or mm39) + virus genome (KT899744.1, NC_009333.1, MH636806.1) + ERCC spike-in controls^44^. Details on mapping assemblies are included below. RNA STAR mapping parameters are as follows: --outSJfilterOverhangMin 15 15 15 15, --outFilterType BySJout, --outFilterMultimapNmax 20, --outFilterScoreMin 1, --outFilterMatchNmin 1, -- outFilterMismatchNmax 2, --outFilterMismatchNoverLmax 0.3, --outFilterIntronMotifs None, --alignIntronMin 20, --alignIntronMax 2000000, --alignMatesGapMax 2000000, -- alignTranscriptsPerReadNmax 20000, --alignSJoverhangMin 15, -- alignSJDBoverhangMin 15, --alignEndsProtrude 10 ConcordantPair, --chimSegmentMin 15, --chimScoreMin 15, --chimScoreJunctionNonGTAG 0 –chimJunctionOverhangMin 18, --chimMultimapNmax 10. STAR GeneCount (per gene read counts) files were used for transcript quantitation. 4sU-Seq data was normalized as reads per million total reads per kilobase pair (RPKM). For all other data, ERCC reads were used to generate standard curves similar to^90^, using their known relative concentrations. All biological replicates had ERCC derived standard curves with R^2^>0.9. ERCC normalized gene counts were calculated as follows:

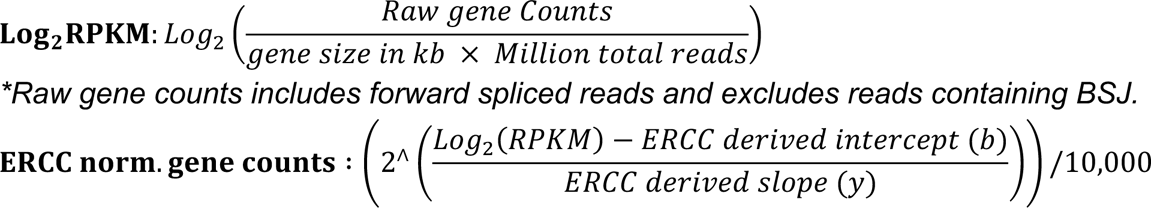

#### CircRNA quantitation

CHARLIE was used to map, annotate, and filter high confidence BSJ as described above. BSJ were quantified from STAR output bams and normalized as RPKM (4sU-Seq) or relative to ERCC spike in controls (all other data). Gene length for circRNA was treated as 0.15 kb as that is the total read length and full circRNA size is unknown.

#### Circ/Linear Transcript Quantitation

High confidence BSJ identified by CHARLIE were quantified from STAR output bams for circRNA and colinear transcripts. CircRNA reads were those which spanned BSJ. Linear reads were those which spanned FSJ in addition to all non-chimeric reads. Circ to linear ratios are the raw reads for circ over linear.

### Flanking cis-element analysis

#### Splice Donor-Acceptor frequency

The 2 nucleotides immediately external to the BSJ were determined for all high confidence circRNAs. For frequency calculations, the splice DA sequences was multiplied by the abundance of a given circRNA in the sample. Thus, circRNA abundance affects total frequency calculations.

#### GC content

The 100 nucleotides (100 nt 5’ + 100 nt 3’) immediately external to the BSJ were determined for all high confidence circRNAs. Flanking sequences were multiplied by the abundance of a given circRNA in the sample. GC content was determined using geecee (Galaxy Version 5.0.0)^91^. To calculate viral gene theoretical GC content, we used the CDS coordinates of KT899744.1 (HSV-1), NC_009333.1 (KSHV), and MH636806.1 (MHV68). To calculate host gene theoretical GC content, we used the gene coordinates of hg38 gencode.v36 and mm39 gencode.vM29^92^.

#### Motif enrichment and RBP predictions

The 100 nucleotides (100 nt 5’ + 100 nt 3’) immediately external to the BSJ were determined for all high confidence circRNAs. Motif discovery and comparison were performed using MEME suite^93^. Motif discovery was performed using MEME v5.5.4 with the following parameters: -oc. -nostatus -time 14400 -mod anr -nmotifs 5 -minw 6 -maxw 10 -objfun classic -minsites 2 -revcomp - markov_order 1. Only motifs with p-values <0.05 were reported. Predicted RBP partners were determined using Tomtom v5.5.4 with the following parameters: -no-ssc -oc. - verbosity 1 -min-overlap 5 -mi 1 -dist pearson -evalue -thresh 10.0 -time 300. Motifs were searched using their species appropriate CISBP-RNA homo sapien or mus musculus database^94^. Only RBP with p-values <0.05 were reported.

### Genome assemblies

<u>HSV-1</u>: KT899744.1, coding sequence (CDS) annotation used for transcript quantification

<u>KSHV</u>: NC_009333.1, CDS annotation used for transcript quantification

<u>MHV68</u>: MH636806.1^95^ modified to remove the beta-lactamase gene (Δ103,908-105,091), CDS annotation used for transcript quantification

<u>Human</u>: hg38, gencode.v36^92^

<u>Mouse</u>: mm39, gencode.vM29^92^

<u>ERCC Spike-In:</u> available from ThermoFisher (#4456740)

## DATA AVAILABILITY

<u>HSV-1 infection:</u> SRR19779319, SRR19779318, SRR19779317, SRR19779316, SRR19779315, SRR19779314, SRR19779313, SRR19779311

<u>HSV-1 infection with spliceosome inhibition:</u> SRR19787559, SRR19787558, SRR19787557, SRR19787556, SRR19787555

<u>HSV-1 murine infection:</u> SRR19792335, SRR19792334, SRR25824398, SRR25824397, SRR25824395, SRR25824394, SRR25824396

*<u>KSHV infection:</u>* SRR20020770, SRR20020769, SRR20020764, SRR20020763, SRR20020762, SRR25816556, SRR20020765, SRR20020766, SRR20020767, SRR20020768, SRR20020757, SRR20020758, SRR20020759, SRR20020760, SRR20020761

<u>KSHV lytic reactivation with spliceosome inhibition:</u> SRR19793315, SRR19793317, SRR19793314

<u>4sU-Seq of KSHV reactivation:</u> SRR19792341, SRR19792340, SRR19792338, SRR19792337, SRR19792339, SRR19792336

<u>ORF57 eCLIP:</u> SRR19793302, SRR19793303

<u>MHV68 infection:</u> SRR19792326, SRR19792325, SRR19792322, SRR19792323, SRR19792324, SRR19792321

<u>MHV68 murine infection:</u> GSE227764

## ACKNOWLEDGEMENTS

We would like to thank Neal DeLuca and Takanobu Tagawa for thoughtful discussions related to this article. Additional thanks to the National Cancer Institute CCR Genomics Core and CCR Illumina Sequencing facility for technical support. The resources of the NIH High-Performance Computing Biowulf Cluster were utilized for all computational needs. This work was supported with funds from the National Institutes of Health, Division of Intramural Research NIAID, T.M.K., ZIA AI000712; NCI, L.T.K., ZIA BC011953; NCI, J.M.Z., ZIA BC011176. Additional grant support R01-AI123165 to N.K.C.

## AUTHOR CONTRIBUTIONS

The conceptualization of this study was done by S.E.D. and J.M.Z. Data curation was performed by S.E.D and V.N.K. Funds to conduct the study were provided by T.M.K., L.T.K., N.K.C., J.M.Z. The experiments in this study were performed by S.E.D., J.H.A., T.M.K., C.H.H, L.T.K., and N.K.C. Methodology was provided by S.E.D., J.H.A., L.T.K., and N.K.C. The original draft of the manuscript was composed by S.E.D. Subsequent writing, review and editing of the manuscript was performed by S.E.D., V.N.K., J.H.A., T.M.K., L.T.K., N.K.C., and J.M.Z.

