## Supplementary Information for "Noncanonical circRNA biogenesis driven by alpha and gamma herpesviruses"

### Table of Contents

Description of Supplementary Datasets
Supplementary Methods
Supplementary Tables
- Table 1. Summary of RNA-Seq data from lytic, latent, and reactivation models.
- Table 2. High confidence circRNA calls in lytic, latent, and reactivation models.
- Table 3. Viruses and cells used in study
- Table 4. Oligos used in study

Supplementary Figures
- Figure 1-1. Viral gene expression in HSV-1, KSHV, and MHV68 infection models
- Figure 1-2: Reproducibility of high confidence circRNA calls
- Figure 1-3. Size distribution of high confidence viral and host circRNAs
- Figure 1-4. CircRNA RNase R resistance
- Figure 1-5. Low-copy number HSV-1 circRNAs called in murine models
- Figure 1-6. *In silico* circRNA-miRNA-mRNA interaction networks
- Figure 2-1. Prominent examples of herpesvirus circRNAs
- Figure 2-2. Herpesvirus circRNA copy levels during lytic infection
- Figure 3-1. CircRNA cis-element analysis for additional lytic infection models
- Figure 3-2. Protein levels of predicted RBP-circRNA partners during infection
- Figure 4-1. Impact of spliceosome inhibition on infection models
- Figure 4-2. Impact of spliceosome depletion on KSHV circRNA levels.
- Figure 5-1. HSV-1 and KSHV gene expression in wildtype and perturbed models
- Figure 5-2. MHV68 gene expression in wildtype and perturbed models
- Figure 5-3. HSV-1 and KSHV transcript profiles in wildtype and perturbed models
- Figure 6-1. ORF57 eCLIP heatmap
- Figure 6-2. ORF57 regulation of linear viral transcripts
- Figure 6-3. ORF57 dependence of PAN BSJ variants
- Supplementary References

**Other supporting materials for this manuscript include the following:**

Supplementary Dataset 1

### DESCRIPTION OF SUPPLEMENTARY DATASETS

**Supporting Dataset 1. Summary of high confidence viral circRNAs**

Description of all high confidence viral circRNAs identified in Fig. 1 RNA-Seq. CircRNAs were determined using CHARLIE. Genomic positions are relative to the following viral genome assemblies: HSV-1: KT899744.1, KSHV: NC_009333.1; MHV68: MH636806.1 modified to remove the beta-lactamase gene (Δ103,908-105,091). Internal and external sequences are reported for the sense strand. BSJ counts are raw values from individual biological replicates.

### SUPPLEMENTARY METHODS

#### *In silico* prediction of circRNA-miRNA-mRNA networks

Expected full length sequences (sequence between 3' and 5' back splice junctions) for viral latency circRNAs was used as input. Prediction of miRNA binding sites was performed using RNAHybrid, with a Gibbs free energy threshold (circLAT: -30, circK12: -10, circORF66: -25), disallowing G:U in seed, and helix constraint from 2 to 8[1]. miRNA-mRNA interactions were determined using QIAGEN Ingenuity Pathway Analysis (IPA) v.01-22-01 MicroRNA Target Filter. We filtered mRNA targets by those with experimentally observed or high predicted value. mRNA networks were generated using IPA[2] expression analysis. circRNA-miRNA-mRNA network diagrams were made in Cytoscape[3].

#### Absolute (ddPCR) Quantification of Transcripts

Total RNA was isolated from cells using the Direct-zol RNA MiniPrep Kit (Zymo), following manufacturer's instructions. 0.2-1 ug total RNA was reverse transcribed with random decamers using ReverTra Ace-$\alpha$ qPCR RT Master Mix (Toyobo). cDNA was measured with either divergent (circRNA) or convergent (mRNA) primers using the QX200 Droplet Digital PCR System (Bio-Rad).

#### Cell Viability Assays

MRC-5 or iSLK-BAC16 cells were treated with 10, 30, 60 uM Isoginkgetin (CAS #548-19-6, Sigma #416154) or 12.5, 25, 50 nM Pladienolide B (CAS #445493-23-2, SantaCruz #sc-391691). When indicated (vehicle), DMSO (0.1%) was added to media. After 24 hours of drug treatment, CellTiter-Glo 2.0 Reagent (Promega #G9241) was added 1:1 to media. Luminescence was measured using the Modulus II Multiplate reader (Turner Biosystems). All values are the average of at least three biological replicates, and values were calculated relative to the luminescence of the unpaired "vehicle" sample.

#### Assessment of unspliced and spliced host transcripts

Total RNA was isolated from cells using the Direct-zol RNA MiniPrep Kit (Zymo #R2053), following manufacturer's instructions. For detection of unspliced targets (pre-BRD2, pre-DNAJB1, 18S rRNA) RNA was reverse transcribed with random decamers using the ReverTra Ace-$\alpha$ qPCR RT Master Mix (Toyobo #FSQ-201). For detection of spliced targets (mature-BRD2, mature-DNAJB1) RNA was reverse transcribed with polydT oligos (Invitrogen #N8080128) using the High-Capacity cDNA Reverse Transcription Kit (Applied Biosystems #4374966). cDNA was measured using qPCR and primers which bound within intronic regions (premature) or spanned exon to exon boundaries (mature). We adapted a previously published method, for gel-based detection of spliced and unspliced DNAJB1 [4]. Total RNA was reverse transcribed using the ReverTra Ace-$\alpha$ qPCR RT Master Mix and random decamers. 20 ng cDNA was PCR amplified using 2x KOD One PCR Master Mix (Toyobo # KMM-101) and amplified with the following cycling conditions: 94$^{o}$C 5 min, (94$^{o}$C 30 sec, 60$^{o}$C 30 sec, 68$^{o}$C 2 min) x 35 cycles, 72$^{o}$C 5 min. PCR material was purified using the Qiagen MinElute kit, run on a 2% agarose 1x TBE gel, and visualized with ethidium bromide staining.

#### siRNA depletion in KSHV reactivation model

Subconfluent dishes of iSLK-BAC16 were incubated in OptiMEM (ThermoFisher #31985062) with 50 nM siRNA and Lipofectamine RNAiMAX Transfection Reagent (ThermoFisher #13778150). siRNAs were ON-TARGETplus SMARTpool siRNAs for Human PRPF8 (10594) Horizon #L-012252 or Non-targeting Horizon #D-001810-10-05. After incubation at 37$^{o}$C for 8 hours transfection media was removed and replaced with 1x DMEM 2% Tet-approved FBS. To induce lytic reactivation media containing 1 mM sodium butyrate 1 ug/mL doxycycline was added at the time transfection media was removed. At 72 hours post transfection total RNA was isolated from cells using the Direct-zol RNA MiniPrep Kit (Zymo #R2053), following manufacturer's instructions. RNA was reverse transcribed with random decamers using the ReverTra Ace-$\alpha$ qPCR RT Master Mix (Toyobo #FSQ-201). qPCR was used to measure PRPF8 (TaqMan gene expression assay, ThermoFisher #4331182), RPS13 (Taqman gene expression assay, ThermoFisher ##4351370), premature DNAJB1, mature DNAJB1, premature K15, and mature K15. ddPCR was used to measure circular (divergent primers) and linear (convergent primers) species.

### SUPPLEMENTARY TABLES

**Supplementary Table 1. Summary of RNA-Seq data from lytic, latent, and reactivation models.** Total mapped reads (MR) reported for all reads mapped to the indicated assembly. Percent total (% Total) is assembly-specific mapped reads relative to input reads. Rows marked with “N/A” or not applicable did not have ERCC synthetic spike-in controls added prior to sequencing.

| Virus | Model | Sample | # Replicates | Total Reads | Total mapped reads (MR) | | | % Total | | |
|---|---|---|---|---|---|---|---|---|---|---|
| | | | | | Host | ERCC | Virus | Host | ERCC | Virus |
| HSV-1 | Lytic | MRC-5 Uninf. | 4 | 8.9E+07 | 8.8E+07 | 3.5E+05 | 2.3E+01 | 98% | 0.4% | 0% |
| | | MRC-5 12 hpi | 4 | 9.2E+07 | 4.6E+07 | 6.9E+05 | 4.3E+07 | 50% | 0.7% | 47% |
| | | MRC-5 24 hpi | 2 | 5.0E+07 | 1.8E+07 | 2.1E+05 | 2.9E+07 | 36% | 0.4% | 57% |
| | | RNaseR MRC5 12 hpi | 2 | 6.8E+07 | 3.9E+07 | N/A | 2.7E+07 | 58% | N/A | 40% |
| | Latent | Murine TG Uninf. | 4 | 1.2E+08 | 1.1E+08 | 1.9E+05 | 3.0E+00 | 91% | 0.2% | 0.00% |
| | | Murine TG Inf. | 4 | 1.4E+08 | 1.2E+08 | 2.1E+05 | 4.4E+04 | 91% | 0.2% | 0.03% |
| | Explant induced reactivation | Murine TG explant Uninf. | 3 | 8.7E+07 | 8.2E+07 | 8.8E+05 | 0.0E+00 | 95% | 1.0% | 0% |
| | | Murine TG explant Inf. | 3 | 8.8E+07 | 8.3E+07 | 9.8E+05 | 5.2E+04 | 95% | 1.1% | 0.06% |
| | | Murine TG explant Inf. + ACV | 3 | 8.5E+07 | 8.0E+07 | 9.4E+05 | 5.9E+04 | 94% | 1.1% | 0.07% |
| | Drug enhanced reactivation | Murine TG explant Uninf. +JQ1 | 3 | 9.3E+07 | 8.9E+07 | 1.1E+06 | 3.0E+00 | 95% | 1.2% | 0.00% |
| | | Murine TG explant Inf. +JQ1 | 3 | 9.1E+07 | 8.6E+07 | 1.1E+06 | 6.3E+04 | 95% | 1.2% | 0.07% |
| KSHV | Lytic | LEC Uninf. | 2 | 1.5E+08 | 1.4E+08 | 1.7E+06 | 2.0E+01 | 98% | 1.2% | 0.0% |
| | | LEC 3dpi | 2 | 1.5E+08 | 1.1E+08 | 1.4E+06 | 3.2E+07 | 75% | 1.0% | 22% |
| | | LEC +RNaseR | 2 | 5.2E+07 | 4.8E+07 | N/A | 1.9E+06 | 93% | N/A | 4% |
| | Pre-Latent | HUV Uninf. | 3 | 2.3E+08 | 2.2E+08 | 3.1E+06 | 9.0E+00 | 98% | 1.3% | 0.0% |
| | | HUV 3dpi | 3 | 2.2E+08 | 2.1E+08 | 2.7E+06 | 8.6E+05 | 97% | 1.3% | 0.4% |
| | Lytic reactivation | iSLK-BAC16 Unind. | 8 | 2.6E+08 | 2.5E+08 | 1.7E+06 | 1.5E+05 | 97% | 0.7% | 0.06% |
| | | iSLK-BAC16 3dpi | 6 | 1.7E+08 | 5.3E+07 | 1.5E+06 | 1.1E+08 | 31% | 0.9% | 63% |
| MHV68 | Lytic | 3T3 Uninf. | 2 | 6.6E+07 | 6.0E+07 | 1.3E+05 | 5.2E+01 | 91% | 0.2% | 0.0% |
| | | 3T3 18 hpi | 2 | 6.9E+07 | 4.0E+07 | 2.4E+05 | 2.3E+07 | 58% | 0.3% | 33% |
| | Latent | Murine GC B-Cell Uninf. | 3 | 1.1E+08 | 8.6E+07 | N/A | 1.3E+03 | 81% | N/A | 0.0% |
| | | Murine GC B-Cell Inf. | 6 | 2.5E+08 | 2.2E+08 | N/A | 1.4E+05 | 91% | N/A | 0.06% |
| | Lytic reactivation | HERIT Unind. | 2 | 6.3E+07 | 5.6E+07 | 1.4E+05 | 2.9E+04 | 90% | 0.2% | 0.0% |
| | | HERIT 24 hpi | 2 | 7.0E+07 | 5.8E+07 | 1.4E+05 | 3.7E+06 | 82% | 0.2% | 5% |

**Supplementary Table 2. High confidence circRNA calls in lytic, latent, and reactivation models.** Overview of high confidence circRNA calls made using CHARLIE. All calls required ≥3 BSJ counts to be reported. For functional genome size we used 100% of the viral genome size, and 10% of the host genome reported as megabase pairs (Mbp).

| Virus | Model | Sample | # Unique BSJ variants | | # BSJ MR | | Avg Read Count/BSJ | | BSJ variants/Mbp functional Genome | | BSJ MR/All MR | |
|---|---|---|---|---|---|---|---|---|---|---|---|---|
| | | | Host | Virus | Host | Virus | Host | Virus | Host | Virus | Host | Virus |
| HSV-1 | Lytic | MRC-5 Uninf. | 481 | 0 | 3,327 | 0 | 6.9 | N/A | 1.5 | N/A | 0.004% | N/A |
| | | MRC-5 12 hpi | 924 | 218 | 7,501 | 1,797 | 8.1 | 8.2 | 2.9 | 1,434 | 0.02% | 0.004% |
| | | MRC-5 24 hpi | 576 | 223 | 4,784 | 2,308 | 8.3 | 10.3 | 1.8 | 1,467 | 0.03% | 0.008% |
| | | RNaseR MRC5 12 hpi | 14,184 | 4,318 | 161,051 | 31,870 | 11.4 | 7.4 | 44.3 | 28,413 | 0.4% | 0.1% |
| | Latent | Murine TG Uninf. | 441 | 0 | 4,472 | 0 | 10.1 | N/A | 1.7 | N/A | 0.004% | N/A |
| | | Murine TG Inf. | 513 | 0 | 4,439 | 0 | 8.7 | N/A | 2.0 | 0 | 0.004% | N/A |
| | Explant induced reactivation | Murine TG explant Uninf. | 379 | 0 | 3,375 | 0 | 8.9 | N/A | 1.5 | N/A | 0.004% | N/A |
| | | Murine TG explant Inf. | 343 | 0 | 2,946 | 0 | 8.6 | N/A | 1.3 | N/A | 0.004% | N/A |
| | | Murine TG explant Inf. + ACV | 431 | 0 | 2,794 | 0 | 6.5 | N/A | 1.7 | N/A | 0.003% | N/A |
| | Drug enhanced reactivation | Murine TG explant Uninf. +JQ1 | 431 | 0 | 4,228 | 0 | 9.8 | N/A | 1.7 | N/A | 0.005% | N/A |
| | | Murine TG explant Inf. +JQ1 | 418 | 0 | 3,749 | 0 | 9.0 | N/A | 1.6 | 0 | 0.004% | N/A |
| KSHV | Lytic | LEC Uninf. | 3,270 | 0 | 24,024 | 0 | 7.3 | N/A | 10.2 | N/A | 0.02% | N/A |
| | | LEC 3dpi | 2,664 | 630 | 18,482 | 3,200 | 6.9 | 5.1 | 8.3 | 4,566 | 0.02% | 0.01% |
| | | LEC +RNaseR | 4,334 | 907 | 38,944 | 4,809 | 9.0 | 5.3 | 13.5 | 6,574 | 0.08% | 0.3% |
| | Pre-Latent | HUV Uninf. | 2,384 | 0 | 48,097 | 0 | 20.2 | N/A | 7.5 | N/A | 0.02% | N/A |
| | | HUV 3dpi | 5,581 | 12 | 50,053 | 37 | 9.0 | 3.1 | 17.4 | 87 | 0.02% | 0.004% |
| | Lytic reactivation | iSLK-BAC16 Unind. | 1,458 | 0 | 15,447 | 0 | 10.6 | N/A | 4.6 | 0 | 0.006% | N/A |
| | | iSLK-BAC16 3dpi | 748 | 1,578 | 7,294 | 11,212 | 9.8 | 7.1 | 2.3 | 11,437 | 0.01% | 0.01% |
| MHV68 | Lytic | 3T3 Uninf. | 280 | 0 | 1,848 | 0 | 6.6 | N/A | 1.1 | N/A | 0.003% | N/A |
| | | 3T3 18 hpi | 489 | 489 | 3,504 | 2,098 | 7.2 | 4.3 | 1.9 | 4095 | 0.009% | 0.009% |
| | Latent | Murine GC B-Cell Uninf. | 1,474 | 0 | 6,728 | 0 | 4.6 | N/A | 5.7 | N/A | 0.008% | N/A |
| | | Murine GC B-Cell Inf. | 2,883 | 7 | 13,798 | 25 | 4.8 | 3.6 | 11.1 | 59 | 0.006% | 0.02% |
| | Lytic reactivation | HERIT Unind. | 209 | 0 | 1,326 | 0 | 6.3 | N/A | 0.8 | 0 | 0.002% | N/A |
| | | HERIT 24 hpi | 344 | 24 | 2,063 | 100 | 6.0 | 4.2 | 1.3 | 201 | 0.004% | 0.003% |

**Supplementary Table 3. Viruses and cells used in study**

| Virus | Mutant Genotype | Complementing cell line | Herpesvirus |
|---|---|---|---|
| **n199**[5] | ICP22 nonsense | N/A | **HSV-1** |
| **n12**[6] | ICP4 nonsense | E5[7] | |
| **5dl1.2**[8] | ICP27 deletion | E11[9] | |
| **KOS**[10] | Wildtype | N/A | |
| **iSLK-ΔORF57**[11] | ORF57 deletion | N/A | **KSHV** |
| **ORF24stop**[12] | ORF24 nonsense | N/A | |
| **BAC16**[13] | Wildtype | N/A | |
| **ORF50stop**[14] | ORF50 stop | CS-RTA4[15] | **MHV68** |
| **H2B-YFP**[16] | Wildtype BAC-derived reporter virus | N/A | |

### Supplementary Table 4. Oligos used in study

| Type | Name | Sequence (5' to 3') | Target Host | Application |
|---|---|---|---|---|
| N/A | **DNAJB1_ F** | GAACCAAAATCACTTTCCCCAAGGAAGG | Human | PCR |
| N/A | **DNAJB1_R** | AATGAGGTCCCCACGTTTCTCGGGTGT | Human | PCR |
| Convergent | **hu_GAPDH_F** | CAGAACATCATCCCTGCCTCTACT | Human | qPCR |
| Convergent | **hu_GAPDH_R** | GCCGAGCTTCCCGTTCA | Human | qPCR |
| Convergent | **hu_RPS13_F** | TCGGCTTTACCCTATCGACGCAG | Human | qPCR |
| Convergent | **hu_RPS13_R** | ACGTACTTGTGCAACACCATGTGA | Human | qPCR |
| Convergent | **hu_PreDNAJB1_F** | GGCCTGATGGGTCTTATCTATGG | Human | qPCR |
| Convergent | **hu_PreDNAJB1_R** | TTAGATGGAAGCTGGCTCAAGAG | Human | qPCR |
| Convergent | **hu_PreBRD2_F** | AGGTAATGTCACAGGATGGGAAGT | Human | qPCR |
| Convergent | **hu_PreBRD2_R** | CCCTGCTGCCTTTCTCTAACC | Human | qPCR |
| Convergent | **hu_MatureDNAJB1_F** | GAACCAAAATCACTTTCCCCAAGGAAGG | Human | qPCR |
| Convergent | **hu_MatureDNAJB1_R** | AATGAGGTCCCCACGTTTCTCGGGTGT | Human | qPCR |
| Convergent | **hu_MatureBRD2_F** | CAAAATTATAAAACAGCCTATGGACATG | Human | qPCR |
| Convergent | **hu_MatureBRD2_R** | TTTTCCAGCGTTTGTGCCATTAGGA | Human | qPCR |
| Convergent | **18S_F** | GTAACCCGTTGAACCCCATT | Human/Mouse | qPCR |
| Convergent | **18S_R** | CCATCCAATCGGTAGTAGCG | Human/Mouse | qPCR |
| Convergent | **7SK_F** | TAAGAGCTCGGATGTGAGGGCGATCTG | Human/Mouse | qPCR |
| Convergent | **7SK_R** | CGAATTCGGAGCGGTGAGGGAGGAAG | Human/Mouse | qPCR |
| Convergent | **mmuGAPDH_F** | CTGACGTGCCGCCTGGAGAAAC | Mouse | qPCR |
| Convergent | **mmuGAPDH_R** | CCCGGCATCGAAGGTGGAAGAGT | Mouse | qPCR |
| Convergent | **hsv_ICP0_F** | CCCACTATCAGGTACACCAGCTT | HSV-1 | qPCR |
| Convergent | **hsv_ICP0_R** | CTGCGCTGCGACACCTT | HSV-1 | qPCR |
| Convergent | **hsv_UL23_F** | ACCCGCTTAACAGCGTCAACA | HSV-1 | qPCR |
| Convergent | **hsv_UL23_R** | CCAAAGAGGTGCGGGAGTTT | HSV-1 | qPCR |
| Convergent | **hsv_UL29_F** | CGAACTTGCGGGTGCGGTCAAA | HSV-1 | qPCR |
| Convergent | **hsv_UL29_R** | CATGGTCGTGTTGGGGTTGAGCATC | HSV-1 | qPCR |
| Convergent | **hsv_UL42_F** | GTCCCGCCGCTCCAGAC | HSV-1 | qPCR |
| Convergent | **hsv_UL42_R** | CTTGCTTCTCCGGTCGGG | HSV-1 | qPCR |
| Convergent | **hsv_UL44_F** | GTGACGTTTGCCTGGTTCCTGG | HSV-1 | qPCR |
| Convergent | **hsv_UL44_R** | GCACGACTCCTGGGCCGTAACG | HSV-1 | qPCR |
| Convergent | **hsv_PreICP0_F** | GATCCAAAGGACGGACCCAG | HSV-1 | qPCR |
| Convergent | **hsv_PreICP0_R** | GATTTCCCGCGTCAATCAGC | HSV-1 | qPCR |
| Convergent | **hsv_MatureICP0_F** | GCGAGTACCCGCCGGCCTGA | HSV-1 | qPCR |
| Convergent | **hsv_MatureICP0_R** | CTCGAACAGTTCCGTGTCC | HSV-1 | qPCR |
| Convergent | **hsv_PreUL15_F** | CCCACCCACATACACACACA | HSV-1 | qPCR |
| Convergent | **hsv_PreUL15_R** | CTCCTCAAGCGATCCCGAAT | HSV-1 | qPCR |
| Convergent | **hsv_MatureUL15_F** | CCCGAGTGGACCACGTTAAA | HSV-1 | qPCR |
| Convergent | **hsv_MatureUL15_R** | CTCGTCGACAAAGAGCAGGT | HSV-1 | qPCR |
| Convergent | **hsv_LAT_5′exon_F** | TTCGTTTTCCCCGTTTCG | HSV-1 | qPCR |
| Convergent | **hsv_LAT_5'exon_R** | CAGACGGGTTAAAGAAACAGAAACC | HSV-1 | qPCR |
| Convergent | **hsv_LAT_3'exon_F** | CGCCTTCCCGAAGAAACTCA | HSV-1 | qPCR |
| Convergent | **hsv_LAT_3'exon_R** | CGCTCAATGAACCCGCATT | HSV-1 | qPCR |
| Convergent | **hsv_LAT_Intron_F** | TGTGTGGTGCCCGTGTCTT | HSV-1 | qPCR |
| Convergent | **hsv_LAT_Intron_R** | CCAGCCAATCCGTGTCGG | HSV-1 | qPCR |
| Convergent | **kshv_PAN_F** | GCTCGCTGCTTGCCTTCTT | KSHV | qPCR |
| Convergent | **kshv_PAN_R** | CCAAAAGCGACGCAATCAA | KSHV | qPCR |
| Convergent | **kshv_ORF6_F** | CTGCCATAGGAGGGATGTTTG | KSHV | qPCR |
| Convergent | **kshv_ORF6_R** | CCATGAGCATTGCTCTGGCT | KSHV | qPCR |
| Convergent | **kshv_K8.1_F** | CCGTCGGTGTGTAGGGATAAAG | KSHV | qPCR |
| Convergent | **kshv_K8.1_R** | GTCGTTGTAGTGGTGGCAGAAA | KSHV | qPCR |
| Convergent | **kshv_K12_F** | ACCGAGTGCTTTAATGCGGA | KSHV | qPCR |
| Convergent | **kshv_K12_R** | AAGCACAATCACGGTTGCAC | KSHV | qPCR |
| Convergent | **kshv_PreORF57_F** | TCCCATTTCTAACGTATCGTGCT | KSHV | qPCR |
| Convergent | **kshv_PreORF57_R** | CTGTAATATCAAACGCGATAAATGAG | KSHV | qPCR |
| Convergent | **kshv_MatureORF57_F** | CAAGCAATGATAGACATGGACATT | KSHV | qPCR |
| Convergent | **kshv_MatureORF57_R** | GTCCCTCGATTCGTCAAACT | KSHV | qPCR |
| Convergent | **mhv_ORF6_F** | AGGGACAGATTTCCTCAGGTGC | MHV68 | qPCR |
| Convergent | **mhv_ORF6_R** | CTGGCGTGGAAGCTGTTACC | MHV68 | qPCR |
| Convergent | **mhv_ORF26_F** | ACTATCTGAGGAGGTGCAC | MHV68 | qPCR |

| | | | | |
|---|---|---|---|---|
| Convergent | **mhv_ORF26_R** | TTTTCCCCTGGGTCAACAC | MHV68 | qPCR |
| Convergent | **mhv_ORF50_F** | GGCCGCAGACATTTAATGAC | MHV68 | qPCR |
| Convergent | **mhv_ORF50_R** | GCCTCAACTTCTCTGGATATGCC | MHV68 | qPCR |
| Convergent | **mhv_LANA_F** | TGTGTGCCAGAAGCTTGTGT | MHV68 | qPCR |
| Convergent | **mhv_LANA_R** | GCCTTATTTTCCCTTACCAG | MHV68 | qPCR |
| Divergent | **hu_circRELL1_F** | ATGTCTGTTAGTGGGGCTGA | Human | qPCR |
| Divergent | **hu_circRELL1_R** | TATCTGCTACCATCGCCTTT | Human | qPCR |
| Divergent | **hu_circTNPO3_F** | TCGTTCCTTACGAATTGGAG | Human | qPCR |
| Divergent | **hu_circTNPO3_R** | CTGCCGGATCTGTAACAACT | Human | qPCR |
| Divergent | **hu_circZKSCAN1_F** | CCTCGAGCTTTGACCTTCATCACG | Human | qPCR |
| Divergent | **hu_circZKSCAN1 _R** | CTCACCTTTATGTCCTGGGAGGT | Human | qPCR |
| Divergent | **hu_circHIPK3_F** | GTCGGCCAGTCATGTATCAA | Human | qPCR |
| Divergent | **hu_circHIPK3_R** | TGGAATACACAACTGCTTGGC | Human | qPCR |
| Divergent | **mmu_circSTAU2_F** | TTCCGTATCCCTCACAGCTC | Mouse | qPCR |
| Divergent | **mmu_circSTAU2_R** | CAGGCCACTAGATCCAAAGC | Mouse | qPCR |
| Divergent | **mmu_circZFP609_F** | AGGAAGGGGAGAATGAGTGC | Mouse | qPCR |
| Divergent | **mmu_circZFP609_R** | TGCCTCCTTGGTCAGAACAT | Mouse | qPCR |
| Divergent | **hsv_circUL1_F** | TAAAGGGCCAAGCGCGTTTC | HSV-1 | qPCR |
| Divergent | **hsv_circUL1_R** | TGGACAGTCGCAAGCAGG | HSV-1 | qPCR |
| Divergent | **hsv_circUL2_F** | CATCGTTAGAGGCGCCGGGAGTG | HSV-1 | qPCR |
| Divergent | **hsv_circUL2_R** | GATGAGCGGCCACGGTTGCCTG | HSV-1 | qPCR |
| Divergent | **hsv_circUL4-5_F** | GGTGATTATCGACTGTCGCGCCG | HSV-1 | qPCR |
| Divergent | **hsv_circUL4-5_R** | GTGTGTCAGCCGTCGGTATTCGTCA | HSV-1 | qPCR |
| Divergent | **hsv_circUL4-8_F** | GGGACGGCCGAGTTTCTC | HSV-1 | qPCR |
| Divergent | **hsv_circUL4-8_R** | CAGTCTTGCTAGGCCCGTC | HSV-1 | qPCR |
| Divergent | **hsv_circUL6-9_F** | GTAAGTGGTGGCGTTGAGGA | HSV-1 | qPCR |
| Divergent | **hsv_circUL6-9_R** | GCTTCCTCCGAGAGCAGAAG | HSV-1 | qPCR |
| Divergent | **hsv_circUL10-12_F** | ATCGCGTAGGGGTCTTCCGC | HSV-1 | qPCR |
| Divergent | **hsv_circUL10-12_R** | CTGCCGATAAACGTCACCAGATGCG | HSV-1 | qPCR |
| Divergent | **hsv_circUL13_F** | CGCACCGTGACTAAGCGTTCC | HSV-1 | qPCR |
| Divergent | **hsv_circUL13_R** | GTGTGGGTTGGAGTGTATGTAGGATGC | HSV-1 | qPCR |
| Divergent | **hsv_circUL29a_F** | CCCGCAGTGGTTCTGGAC | HSV-1 | qPCR |
| Divergent | **hsv_circUL29a_R** | CTGCATGATGGTCCGGGC | HSV-1 | qPCR |
| Divergent | **hsv_circUL29b_F** | TACCCGTGTATCCGTTGCAGC | HSV-1 | qPCR |
| Divergent | **hsv_circUL29b_R** | GTTTGGGGCCTGGGTGCTG | HSV-1 | qPCR |
| Divergent | **hsv_circUL36_F** | CGCTGGTGGCCAGCTC | HSV-1 | qPCR |
| Divergent | **hsv_circUL36_R** | GTCGCTGAGGGCCAGCTG | HSV-1 | qPCR |
| Divergent | **hsv_circUL44a_F** | GAGACTGTGGTGAACCCGTC | HSV-1 | qPCR |
| Divergent | **hsv_circUL44a_R** | GTCCCCCCGCGGACCTTCAC | HSV-1 | qPCR |
| Divergent | **hsv_circUL44b_F** | CTGGTGACTGCCGTGGTG | HSV-1 | qPCR |
| Divergent | **hsv_circUL44b_R** | GAGCGGCAGGTGATCGAG | HSV-1 | qPCR |
| Divergent | **hsv_circUL47_F** | CCGTAGCCAGTCGGTGC | HSV-1 | qPCR |
| Divergent | **hsv_circUL47_R** | CCCGCCATCTCCTCCAGA | HSV-1 | qPCR |
| Divergent | **hsv_circUS8-9_F** | CTGGAAGCAACGAAACGTCC | HSV-1 | qPCR |
| Divergent | **hsv_circUS8-9_R** | GCAATGTTGTCTCCCGGTTG | HSV-1 | qPCR |
| Divergent | **hsv_circICP0a_F** | GAGGAAGGGAGGGAGGAGG | HSV-1 | qPCR |
| Divergent | **hsv_circICP0a_R** | CCCGGTGTCGTTCAACAAAG | HSV-1 | qPCR |
| Divergent | **hsv_circICP0b_F** | CCCGTGGTCCGGGGAG | HSV-1 | qPCR |
| Divergent | **hsv_circICP0b_R** | GCCCCGATGTTCCCCGTC | HSV-1 | qPCR |
| Divergent | **hsv_circLAT_F** | GGGCTGGTGTGCTGTAACA | HSV-1 | qPCR |
| Divergent | **hsv_circLAT_R** | GGGTCGCCATGTTTCCCG | HSV-1 | qPCR |
| Divergent | **kshv_circK2a,b,c,d_F** | ATCCCAGACGTGACTCCTGA | KSHV | qPCR |
| Divergent | **kshv_circK2a_R** | TCCCTAATAGACCGCGGTCA | KSHV | qPCR |
| Divergent | **kshv_circK2b_R** | TTTGGGTGGACTGTAGTGCG | KSHV | qPCR |
| Divergent | **kshv_circK2c_R** | AGAAGCTCCATGACGTCCAC | KSHV | qPCR |
| Divergent | **kshv_circK2d_R** | GCCATCGGCGAGCTTTTTAA | KSHV | qPCR |
| Divergent | **kshv_circK4a_F** | GCAAGCCGGGTGTGATATT | KSHV | qPCR |
| Divergent | **kshv_circK4a_R** | TAACTCCCCTCGTGTGTCCT | KSHV | qPCR |
| Divergent | **kshv_circK4b_F** | TTACCAGTCACTGCTCGCTG | KSHV | qPCR |
| Divergent | **kshv_circK4b_R** | AACGGCGATGTTAGGTCAGG | KSHV | qPCR |
| Divergent | **kshv_circK5_F** | ACCTGTGTAAATTTCGGTGTGGTTT | KSHV | qPCR |
| Divergent | **kshv_circK5_R** | GTCCACTTTTCCCCGGCTAT | KSHV | qPCR |

| | | | | |
|---|---|---|---|---|
| Divergent | **kshv_circPANa,b_F** | GCCCGATTTACACTCAATCCG | KSHV | qPCR |
| Divergent | **kshv_circPANa_R** | CGATTTGAATGACATAGGCGACA | KSHV | qPCR |
| Divergent | **kshv_circPANb_R** | TGGTGCGTTGTGAAGCATTT | KSHV | qPCR |
| Divergent | **kshv_circPANc_F** | ACCAGACGGCAAGGTTTTTA | KSHV | qPCR |
| Divergent | **kshv_circPANc_R** | TCGTTAGTCAACCTAGCAAAACA | KSHV | qPCR |
| Divergent | **kshv_circPANd_F** | TGTTACACAACGCTTTCACCT | KSHV | qPCR |
| Divergent | **kshv_circPANd_R** | TGACAATTTAACGTGCCTAGAGC | KSHV | qPCR |
| Divergent | **kshv_circPAN_ 28646-29488_F** | CTTCAAGCTGACCCTTAAT | KSHV | qPCR |
| Divergent | **kshv_circPAN_ 28646-29488_R** | CAGGCTAGTGCTGTAAT | KSHV | qPCR |
| TAQ Probe | **kshv_circPAN_ 28646-29488_Probe** | /56-FAM/TAAGCAAGT/ZEN/CGAT TTGAATATTTGGTTTCC/3IABKFQ/ | KSHV | qPCR |
| Divergent | **kshv_circORF32-34_F** | CAACCAAAAGGCAGAGTCGT | KSHV | qPCR |
| Divergent | **kshv_circORF32-34_R** | GGCTGAACCCAAGAACTTCA | KSHV | qPCR |
| Divergent | **kshv_circORF45_F** | AAACCGGTAGCAGTGGTAGC | KSHV | qPCR |
| Divergent | **kshv_circORF45_R** | GAGGCGCTCCTTCAATTGGA | KSHV | qPCR |
| Divergent | **kshv_circORF46_F** | CGGTTCCAGGGCTGTAATCA | KSHV | qPCR |
| Divergent | **kshv_circORF46_R** | TGTGCGTTTATGAGCGGAGT | KSHV | qPCR |
| Divergent | **kshv_circORF46-47_F** | TGGTTGCAACAGACGGTCTT | KSHV | qPCR |
| Divergent | **kshv_circORF46-47_R** | TCTCCGGCTGCTGCTTTTAG | KSHV | qPCR |
| Divergent | **kshv_circORF58a_F** | TGGTATTGGCCCTATGCACG | KSHV | qPCR |
| Divergent | **kshv_circORF58a,b,c_R** | GCGCAAACTACAGCAGCAAA | KSHV | qPCR |
| Divergent | **kshv_circORF58b_F** | TGGGTTACTGTCATGCGTGG | KSHV | qPCR |
| Divergent | **kshv_circORF58c_F** | ATGCCCGGGAAGTACATCAC | KSHV | qPCR |
| Divergent | **kshv_circORF59a_F** | GACAGCACAAAGAGGCCTCA | KSHV | qPCR |
| Divergent | **kshv_circORF59a,b_R** | GGAAATGGTGGTCCTGACGA | KSHV | qPCR |
| Divergent | **kshv_circORF59b_F** | TGTGTAAAGTCCCGGGTTGG | KSHV | qPCR |
| Divergent | **kshv_circORF60-62_F** | GGAAGAAGCTCATGGACTGG | KSHV | qPCR |
| Divergent | **kshv_circORF60-62_R** | GACCTAAAAACCCGGAGGAG | KSHV | qPCR |
| Divergent | **kshv_circvIRF4_F** | CTCCGTGTGGATACCAGTGA | KSHV | qPCR |
| Divergent | **kshv_circvIRF4_R** | TGGTTCCACGCAACAGTCT | KSHV | qPCR |
| Divergent | **kshv_circK12a_F** | CATGGCAGTACATTGCAGCG | KSHV | qPCR |
| Divergent | **kshv_circK12a_R** | CCGAAGTCAGTGCCACAATT | KSHV | qPCR |
| Divergent | **kshv_circK12b_F** | AGTGAGGAGGGAGGAGGGCA | KSHV | qPCR |
| Divergent | **kshv_circK12b_R** | CGCTCTCCCAAACCACACGAAT | KSHV | qPCR |
| Divergent | **kshv_circORF72_F** | AAGATTAAGGGCCAACGCGA | KSHV | qPCR |
| Divergent | **kshv_circORF72_R** | TAGTTCCTCAGCTGGCAAGC | KSHV | qPCR |
| Divergent | **mhv_circORF6_F** | CTGCTGGTCATGAGATAGGAAATAAACT | MHV68 | qPCR |
| Divergent | **mhv_circORF6_R** | GCTCATGCAGAGAAGATCCAGC | MHV68 | qPCR |
| Divergent | **mhv_circORF11_F** | GTCAGGGTTCTGGGCCAC | MHV68 | qPCR |
| Divergent | **mhv_circORF11_R** | GCCCACAATTTAACCGCCG | MHV68 | qPCR |
| Divergent | **mhv_circORF20_F** | GATATTTAAGAGCCAGCGGGGTC | MHV68 | qPCR |
| Divergent | **mhv_circORF20_R** | TGGTGGTACTGTGCTGGTTG | MHV68 | qPCR |
| Divergent | **mhv_circORF25-26_F** | GCAGTCTTGTTGCCAATCTTTAATACTC | MHV68 | qPCR |
| Divergent | **mhv_circORF25-26_R** | GAGGAGGTGCACATTTGGC | MHV68 | qPCR |
| Divergent | **mhv_circORF47-48_F** | CCCAAGCATTGTTGCCATATTTCTG | MHV68 | qPCR |
| Divergent | **mhv_circORF47-48_R** | CCTTCACGGGTTTCAAGGTCC | MHV68 | qPCR |
| Divergent | **mhv_circORF52_F** | CATGTAAACACACACAGTACATGATATGAA | MHV68 | qPCR |
| Divergent | **mhv_circORF52_R** | CCTGAAGACTTAACCTTTTGTTTAAGTG | MHV68 | qPCR |
| Divergent | **mhv_circORF66a_F** | ATCTATTGTTCATATTGTGGGGCG | MHV68 | qPCR |
| Divergent | **mhv_circORF66a_R** | CAACGCACTATTCGCCACC | MHV68 | qPCR |
| Divergent | **mhv_circORF66b_F** | GTATGTTTTATTGCAGAGATCAAAAGGAG | MHV68 | qPCR |
| Divergent | **mhv_circORF66b_R** | GATCTAGGGCCTTGAGGAAGG | MHV68 | qPCR |
| Divergent | **mhv_circORF68_F** | GGTTTAGTCCCGCGTTAGAGAAG | MHV68 | qPCR |
| Divergent | **mhv_circORF68_R** | CCCCGATCAGGTGGCTC | MHV68 | qPCR |
| Divergent | **mhv_circM7_F** | GTAATACCCTCTACAGTCTCGGCC | MHV68 | qPCR |
| Divergent | **mhv_circM7_R** | GACCTGGGTCTTCTTTTCTTGGT | MHV68 | qPCR |

### SUPPLEMENTARY FIGURES

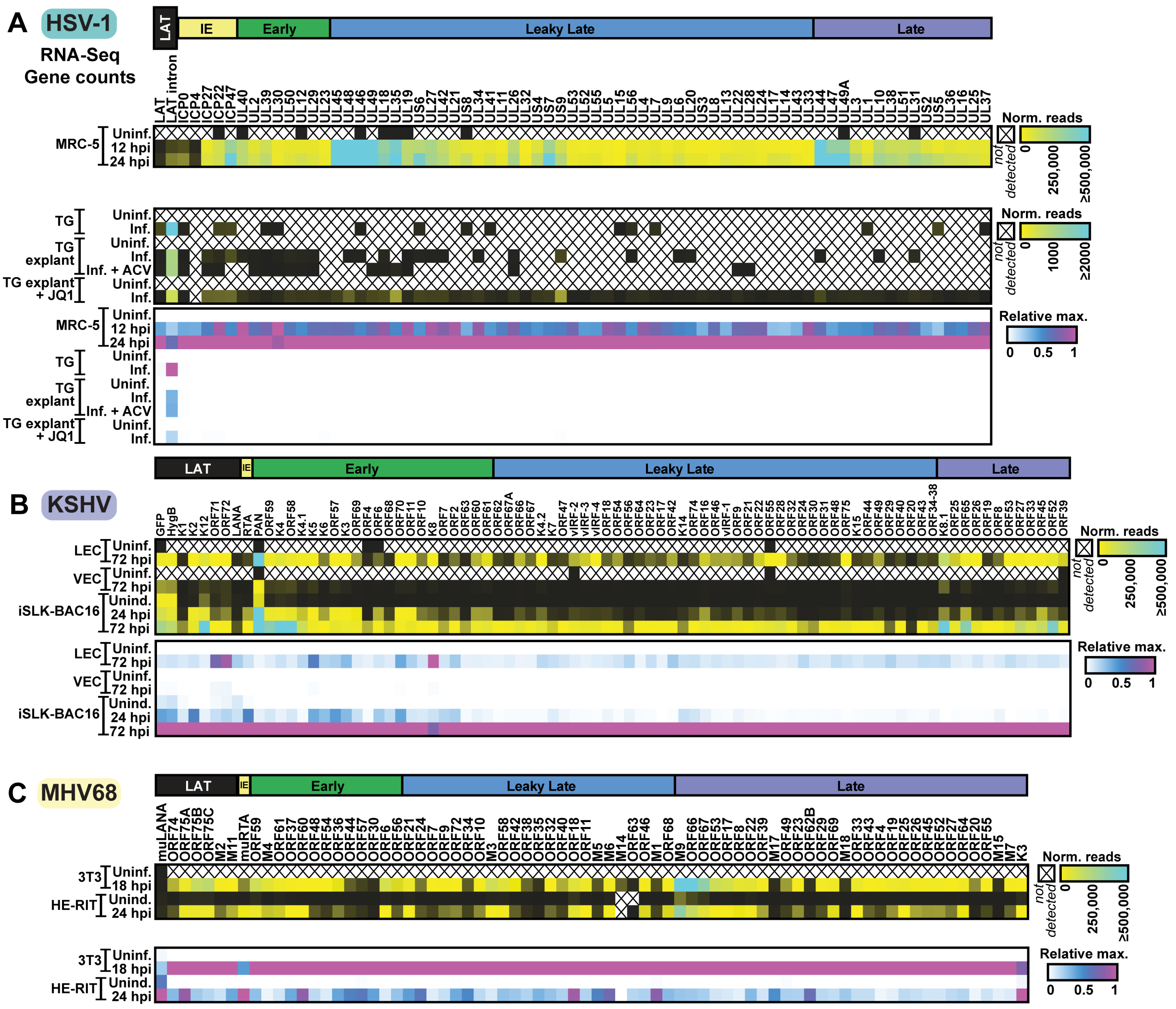

**Supplementary Figure 1-1. Viral gene expression in HSV-1, KSHV, and MHV68 infection models** Viral gene counts for RNA-Seq data in Fig. 1 are plotted as ERCC normalized reads (Norm. reads) or relative maximum in each column (Relative max). Data is the average of biological replicates. Genes are clustered by transcriptional class and labeled as LAT (latent), IE (immediate early), Early, Leaky Late, Late.

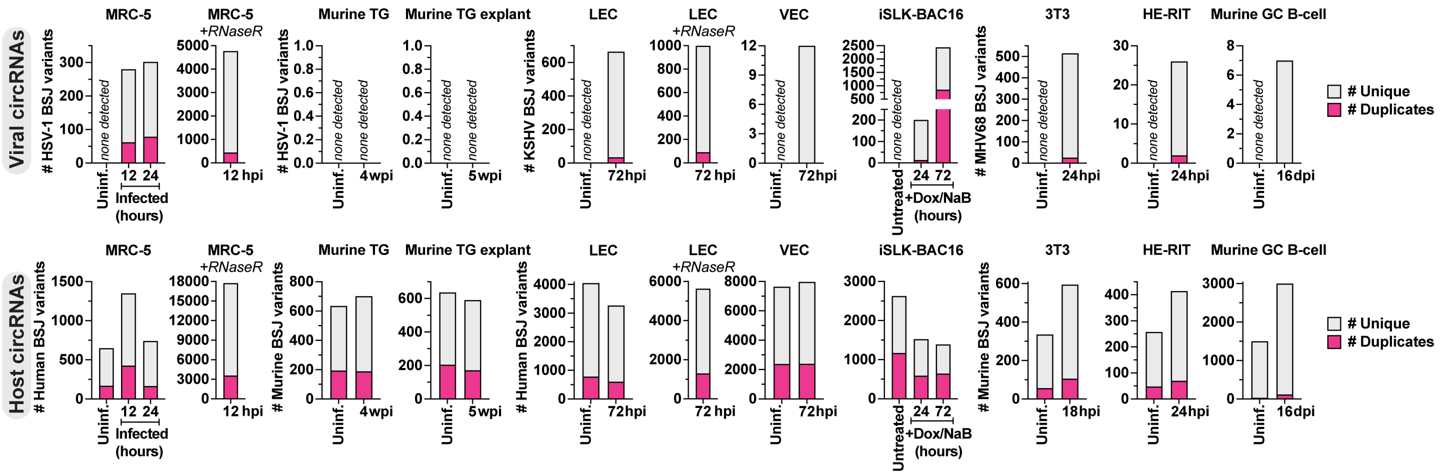


### Supplementary Figure 1-2. Reproducibility of high confidence circRNA calls

High confidence viral and host circRNA calls for data in Fig. 1. The number of unique or duplicate (relative to other biological replicates) high confidence circRNAs are reported.

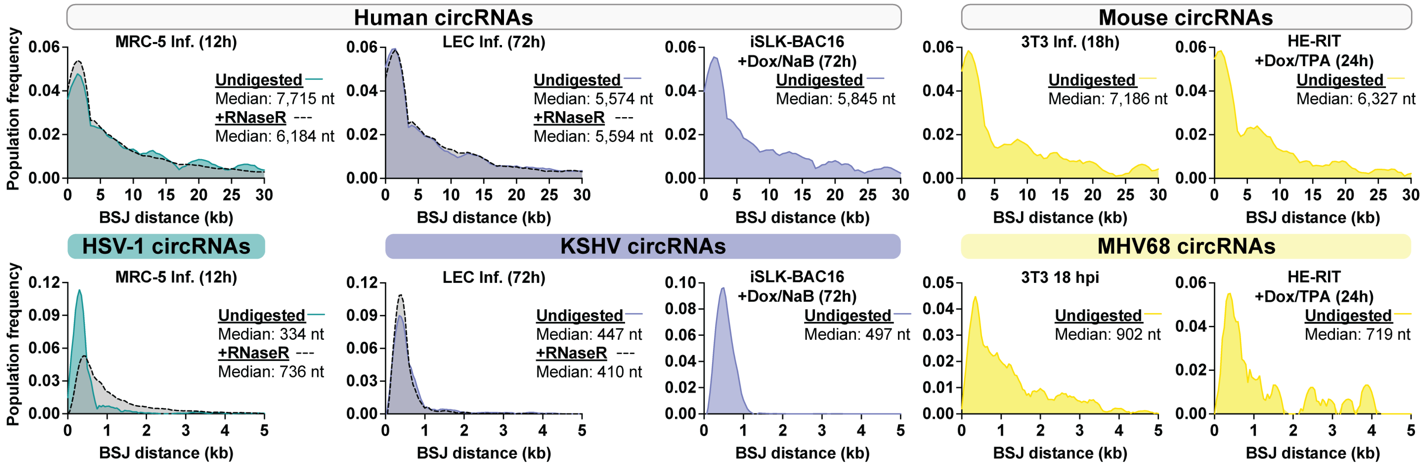


### Supplementary Figure 1-3. Size distribution of high confidence viral and host circRNAs

Population frequency distribution for the expected size (BSJ junction end – BSJ junction start) of circRNAs expressed during lytic infection. Each plot is the sum of all high confidence BSJ calls in the sample. If indicated, samples were treated with RNase R (+RNaseR) prior to sequencing. The median size for a sample is reported in nucleotides (nt).

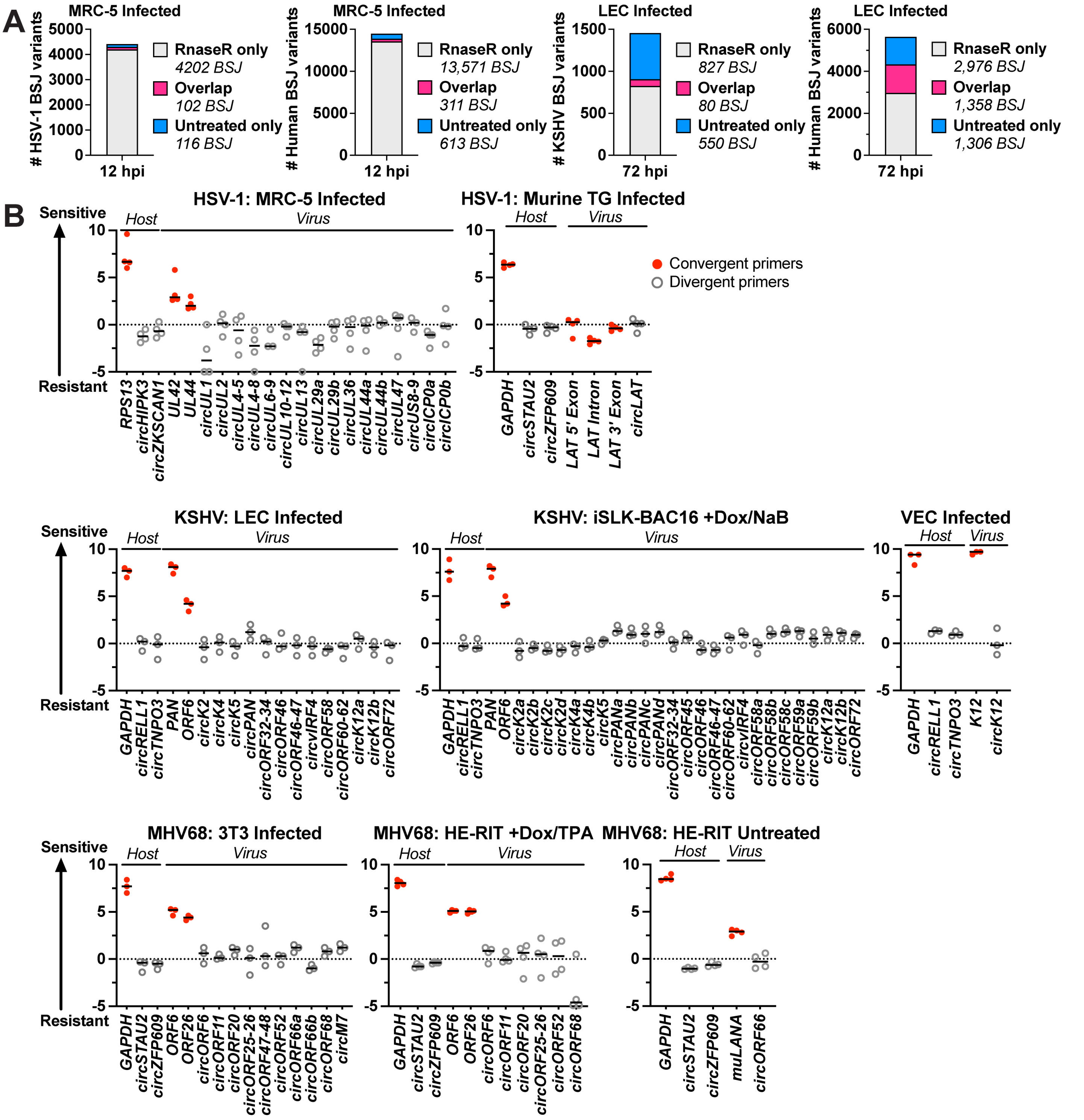


**Supplementary Figure 1-4. CircRNA RNase R resistance**

A) Total RNA was collected from MRC-5 infected with HSV-1 for 12 hours or LEC infected with KSHV for 72 hours. RNA samples were polyA-tailed and RNase R digested (MRC-5) or RNase R digested (LEC). The number of high confidence viral and host BSJ variants found in untreated and/or RNase R treated samples is reported. B) Total RNA was collected from the models described in Fig. 1. and polyA-tailed and RNase R digested (HSV-1), or RNase R digested (KSHV, MHV68). Mock samples were subjected to all steps in parallel except RNase R was not added to the digestion reaction. cDNA samples were assessed by qPCR with divergent (grey dots) or convergent (red dots) primers. Values plotted are delta Ct (RNase R treated sample Ct - Mock treated sample Ct), data points are biological replicates and horizontal lines are the average.

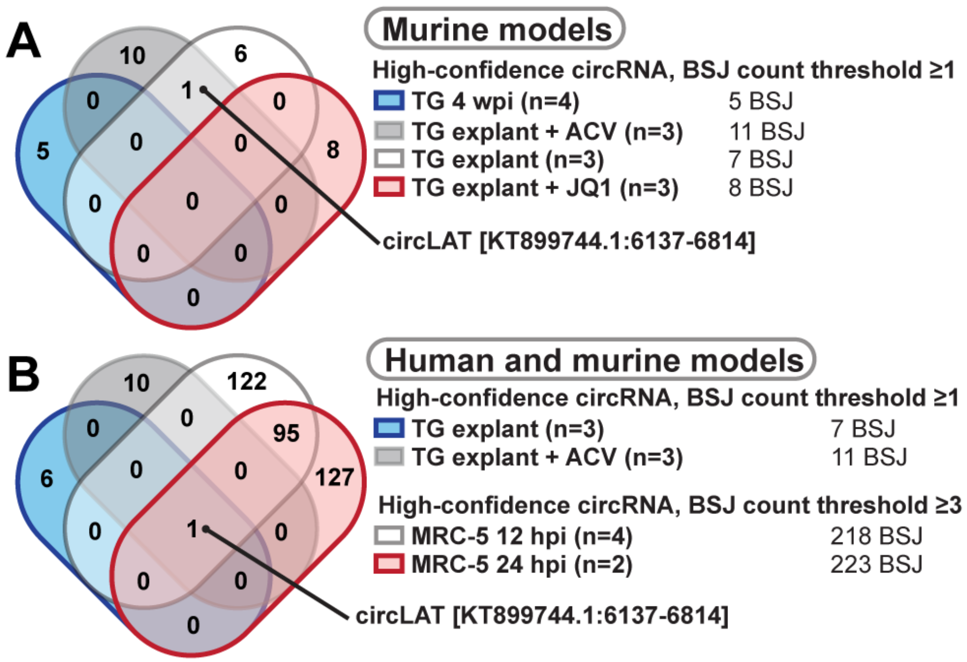


**Supplementary Figure 1-5. Low-copy number HSV-1 circRNAs called in murine models**

Our high confidence circRNA threshold was lowered (count of ≥1 rather than ≥3) to reanalyze murine HSV-1 infection models. The overlapping incidence of these circRNAs is demonstrated for A) murine models and B) murine and human models.

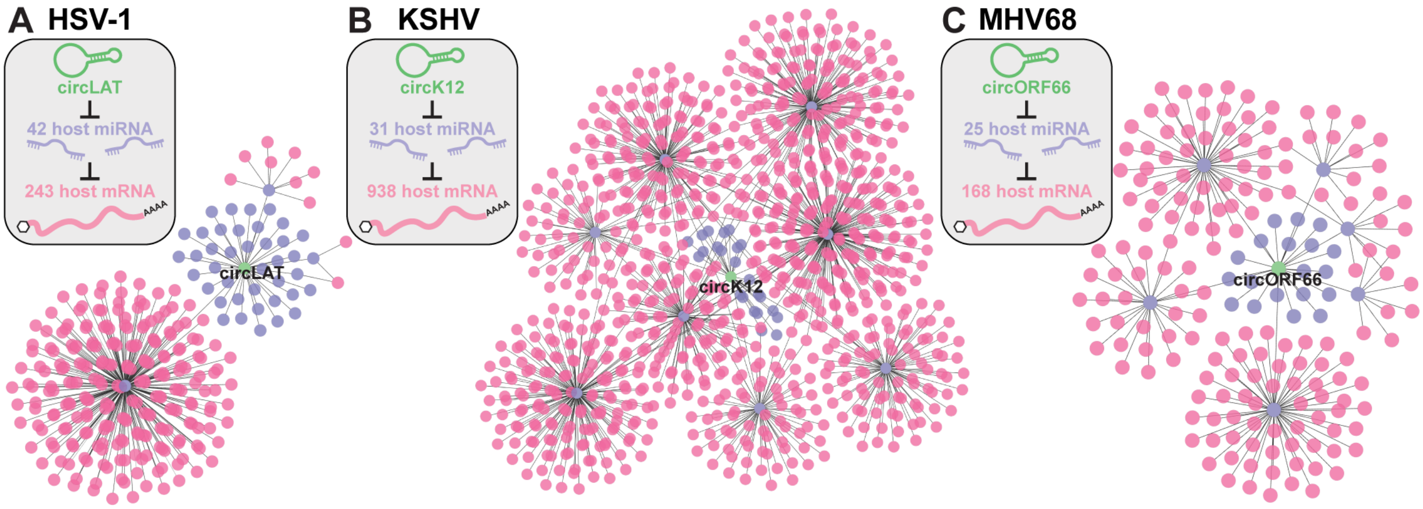


D

**IPA Core Analysis** *(based on predicted mRNA targets)*

| CircRNA | Pathway | p-value |
|---|---|---|
| circLAT | **Aryl Hydrocarbon Receptor Signaling** (*AHR,AHRR,ALDH1A2,ALDH9A1,CDK2,CDK4,CDK6,CYP1B1,NFIC,RARG,RELA,SP1,TRIP11*) | 1.12E-8 |
| | **Senescence Pathway** (*AKT2,ARAF,CDK2,CDK4,CDK6,CHP1,DHCR24,E2F5,ELF4,ELK3,MAPK14,NFATC1,PHF19,RASSF5,SMAD5,SQSTM1*) | 2.24E-7 |
| | **ID1 Signaling Pathway** (*AKT2,ARAF,CAV1,CCN2,CDK4,DVL2,EGR1,GSK3B,MAPK14,PGF,RHOC,SMAD5,STAT3*) | 4.27E-7 |
| circK12 | **CLEAR Signaling Pathway** (*AKT3,ATF2,ATP6AP1,ATP6V0B,ATP6V0E2,ATP6V1E1,ATP6V1E2,ATP6V1F,ATP6V1G2,H2BW2,KRAS,MAP4K3,MAPK11,MAPK13,MAPK6, NTRK3,PIP4P1,PRKAB2,PRKAG1,PRKCD,YWHAE*) | 7.94E-3 |
| | **mTOR Signaling** (*AKT3,EIF3I,EIF4G3,FKBP1A,GPLD1,KRAS,PLD6,PRKAB2,PRKAG1,PRKCD,RHOB,RHOJ,RND3,RPS15A,RPS21*) | 2.63E-2 |
| | **Role of JAK family kinases in IL-6-type Cytokine Signaling** (*ADAM17,BCL2L1,CLCF1,MAPK11,MAPK13,OSMR,PIAS3*) | 4.27E-2 |
| circORF66 | **Integrin and Thrombin Signaling** (*Cdc42,Ppp1r12b*) | 4.37E-2 |

**Supplementary Figure 1-6.** ***In silico*** **circRNA-miRNA-mRNA interaction networks**

A) *In silico* circRNA-miRNA-mRNA interaction networks for circLAT (HSV-1), circK12 (KSHV), and circORF66 (MHV68) variants highlighted in Fig. 1C-D. B) Putative downstream mRNA targets were used to perform over-representation analysis. Pathway hits, mRNA targets present, and p-values are given.

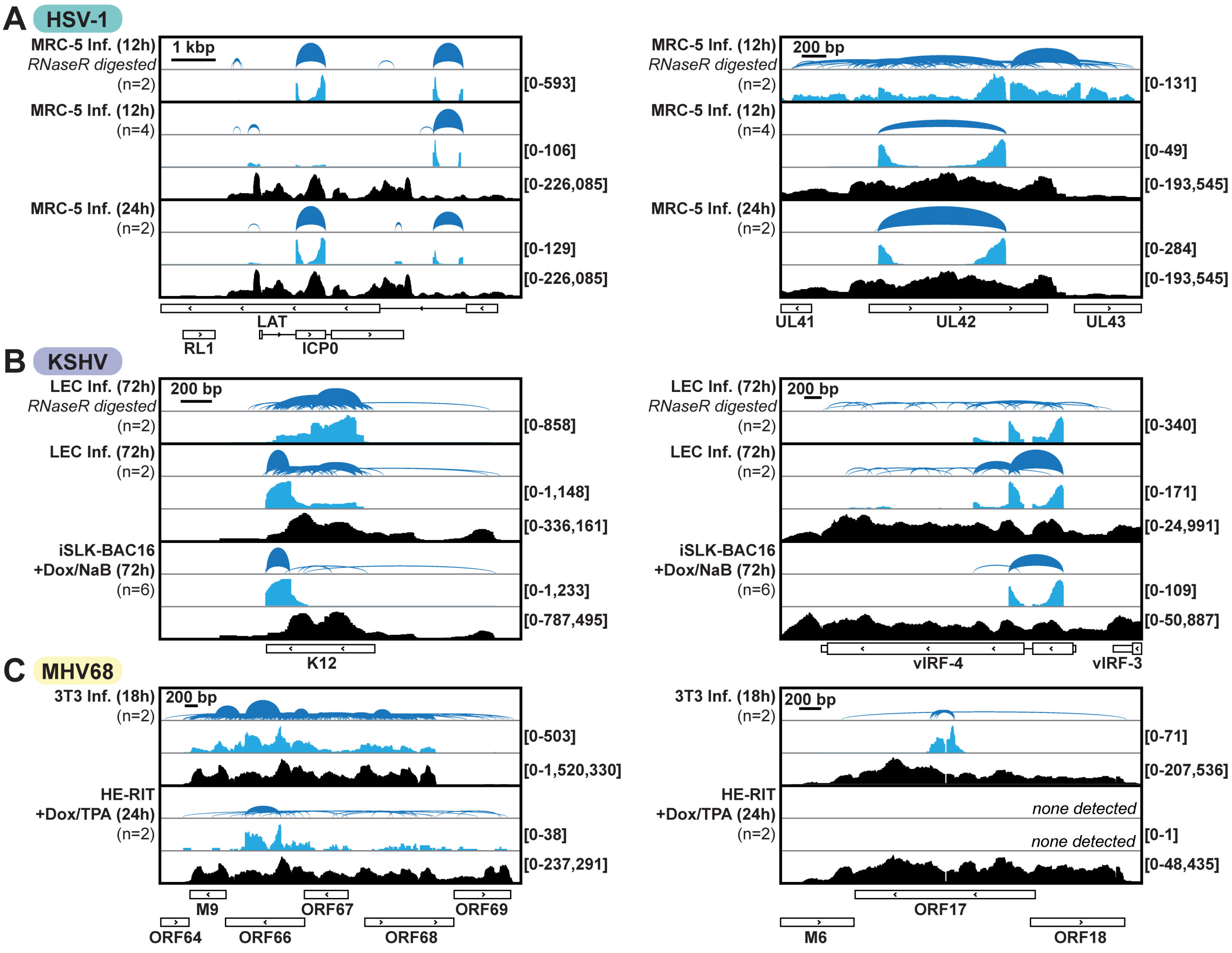


**Supplementary Figure 2-1. Prominent examples of herpesvirus circRNAs**

Visualization of high confidence circRNAs for HSV-1, KSHV, and MHV68 in lytic models described in Fig. 1. If indicated, RNA samples were treated with RNase R (+RNaseR) prior to sequencing. Sashimi plots show high confidence circRNA with arcs proportional to raw BSJ counts. Blue and black traces include circular (back spliced reads) and linear (non-chimeric) reads, respectively. Traces are the sum of raw BSJ or linear read values for all biological replicates. Y-axis minimum and maximum values are shown on the right. Viral genes are shown below.

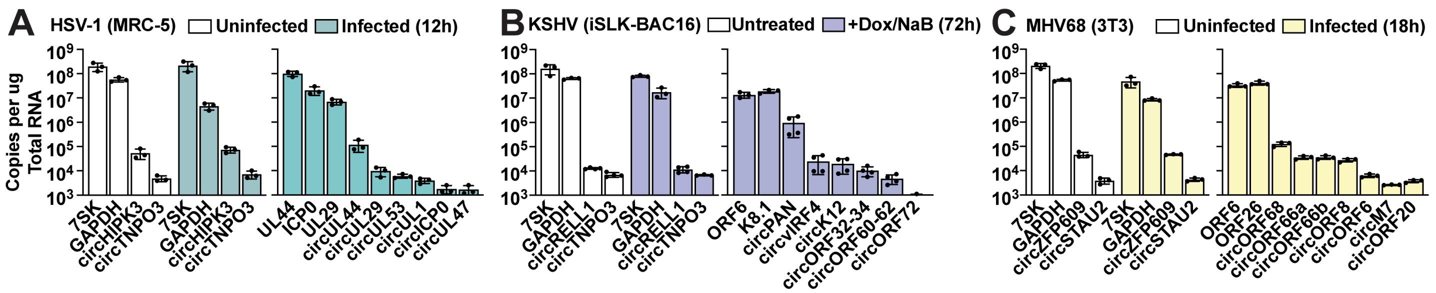


**Supplementary Figure 2-2. Herpesvirus circRNA copy levels during lytic infection**

RNA was collected from A) MRC-5 infected with HSV-1 strain KOS at MOI of 10 PFU/cell for 12 hours, B) LEC infected with KSHV strain BAC16 at MOI of 1 PFU/cell for 72 hours, or C) 3T3 infected with MHV68 strain H2B-YFP at MOI of 5 PFU/cell for 18 hours. cDNA was quantified using digital droplet PCR (ddPCR) and convergent (linear transcripts) or divergent (circular transcripts) primers. Data is normalized to the total amount of RNA in the reverse transcription reaction and plotted as copies per ug total RNA. Data points are biological replicates, column bars are the average, and error bars are standard deviation.

A

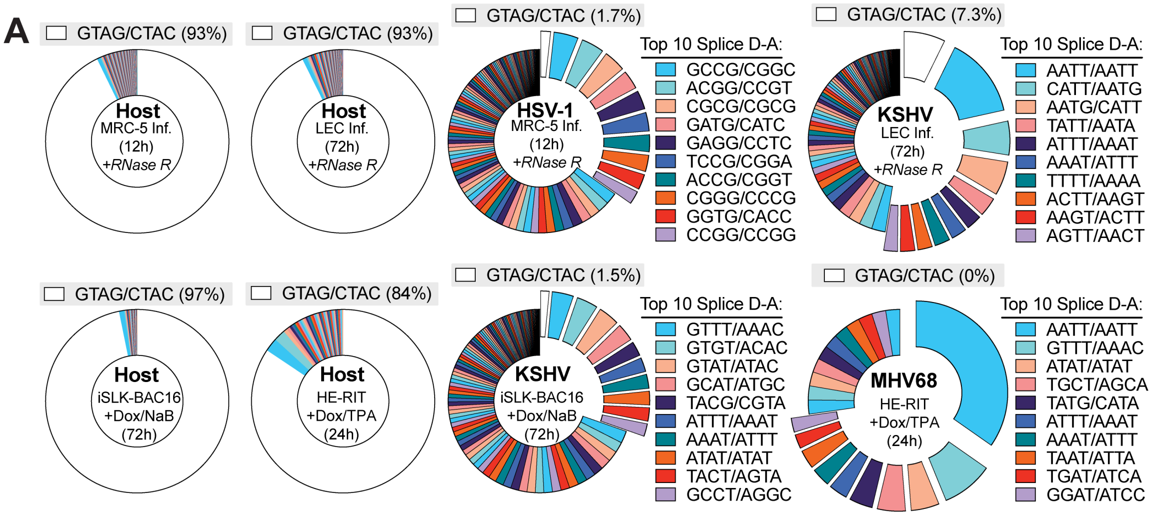


B

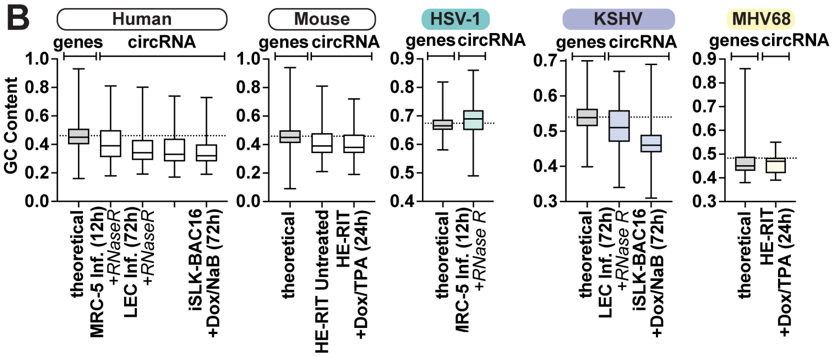


| Sample Group | | | Mean GC content | Discrepancy | P-Value |
|---|---|---|---|---|---|
| | | *Human Genes: theoretical GC usage* | *46.1%* | *N/A* | *N/A* |
| **Human circRNA** | +RNaseR | MRC-5 Inf. (12h) | 41.3% | -4.8% | <0.0001 |
| | +RNaseR | LEC Inf. (72h) | 37.2% | -8.9% | <0.0001 |
| | N/A | iSLK-BAC16 Untreated | 36.4% | -9.7% | <0.0001 |
| | N/A | iSLK-BAC16 +Dox/NaB (72h) | 34.8% | -22.3% | <0.0001 |
| | | *Mouse Genes: theoretical GC usage* | *45.8%* | *N/A* | *N/A* |
| **Mouse circRNA** | N/A | HE-RIT Untreated | 41.2% | -4.6% | <0.0001 |
| | N/A | HE-RIT +Dox/TPA (24h) | 40.2% | -5.6% | <0.0001 |
| | | *HSV-1 Genes: theoretical GC usage* | *67.4%* | *N/A* | *N/A* |
| **HSV-1 circRNA** | +RNaseR | MRC-5 Inf. (12h) | 68.5% | **1.1%** | <0.0001 |
| | | *KSHV Genes: theoretical GC usage* | *53.8%* | *N/A* | *N/A* |
| **KSHV circRNA** | +RNaseR | LEC Inf. (72h) | 51.3% | -2.5% | <0.0001 |
| | N/A | iSLK-BAC16 +Dox/NaB (72h) | 46.4% | -7.4% | <0.0001 |
| | | *MHV68 Genes: theoretical GC usage* | *48.3%* | *N/A* | *N/A* |
| **MHV68 circRNA** | N/A | HE-RIT +Dox/TPA (24h) | 46.1% | -2.2% | <0.0001 |

**Supplementary Figure 3-1. CircRNA cis-element analysis for additional lytic infection models**
Cis-element analysis was performed for high confidence circRNAs identified in RNA-Seq data from Fig. 1. If indicated, samples were treated with RNase R prior to RNA-Seq. A) Splice donor-acceptor frequency for circRNAs identified in lytic infection models, cis-elements are reported as sense and antisense sequences. The percentage of the total which use the canonical splice donor acceptor (GT-AG/CT-AC) are reported above. B) GC content of 100 nucleotides flanking BSJ variants relative to the theoretical GC content of genes for an organism. Wilcoxon t-tests were performed, relative to the theoretical gene GC content, to test significance.

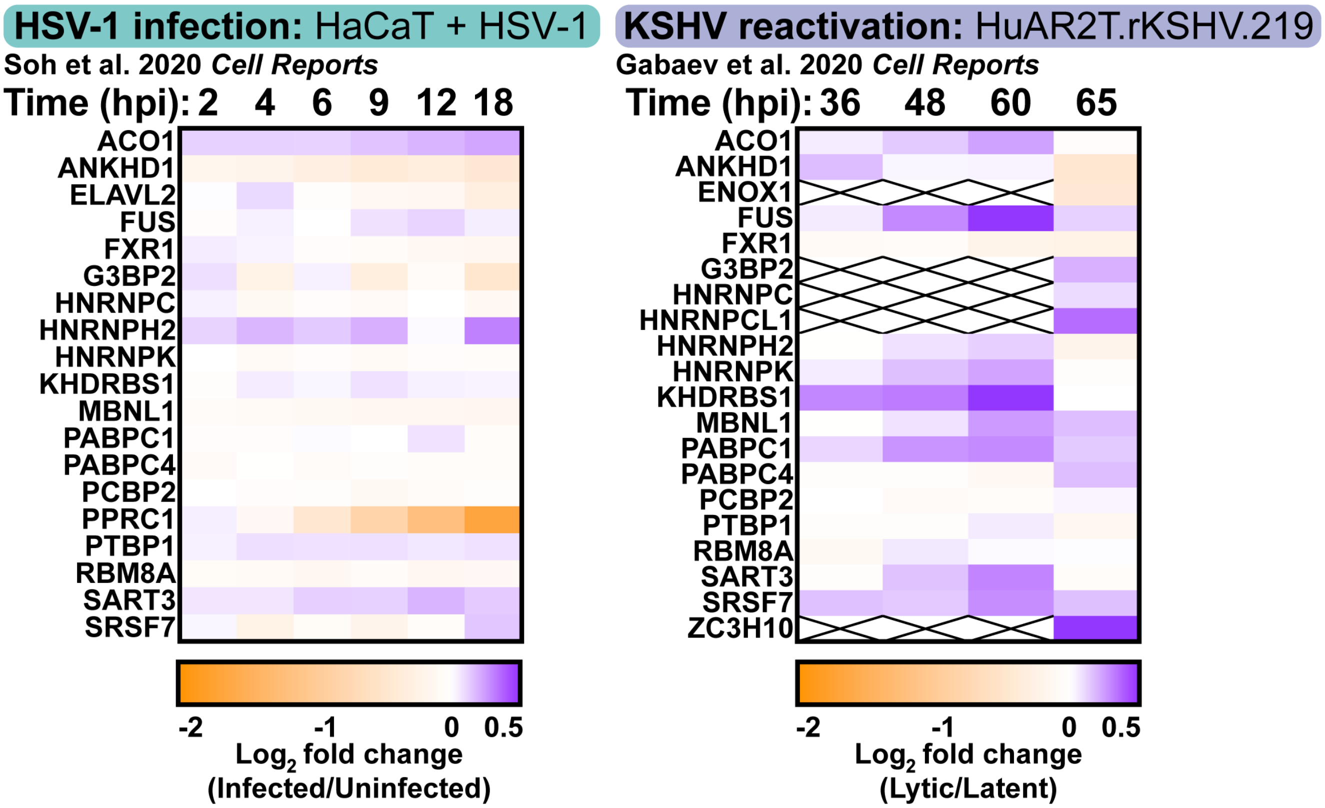


**Supplementary Figure 3-2. Protein levels of predicted RBP-circRNA partners during infection**
Tandem mass tag mass spectrometry data for predicted RBP partners in Fig. 3E from A) HSV-1 infected HaCaT (human immortalized keratinocytes)[17] or B) lytic HuAR2T.rKSHV.219 (immortalized human umbilical vein endothelial cells)[18]. HSV-1 data is the $\log_2$ fold change of infected/mock-infected samples, values are the average of biological duplicates. KSHV data is the $\log_2$ fold change of cells transduced with a lentiviral vector expressing RTA (lytic) verse cells transduced with a lentiviral vector expressing BFP (latent). 36, 48, and 60 hpi data is from one biological replicate. 65 hpi data is the average of biological duplicates. Squares marked with an “X” did not have any peptides detected for the indicated protein.

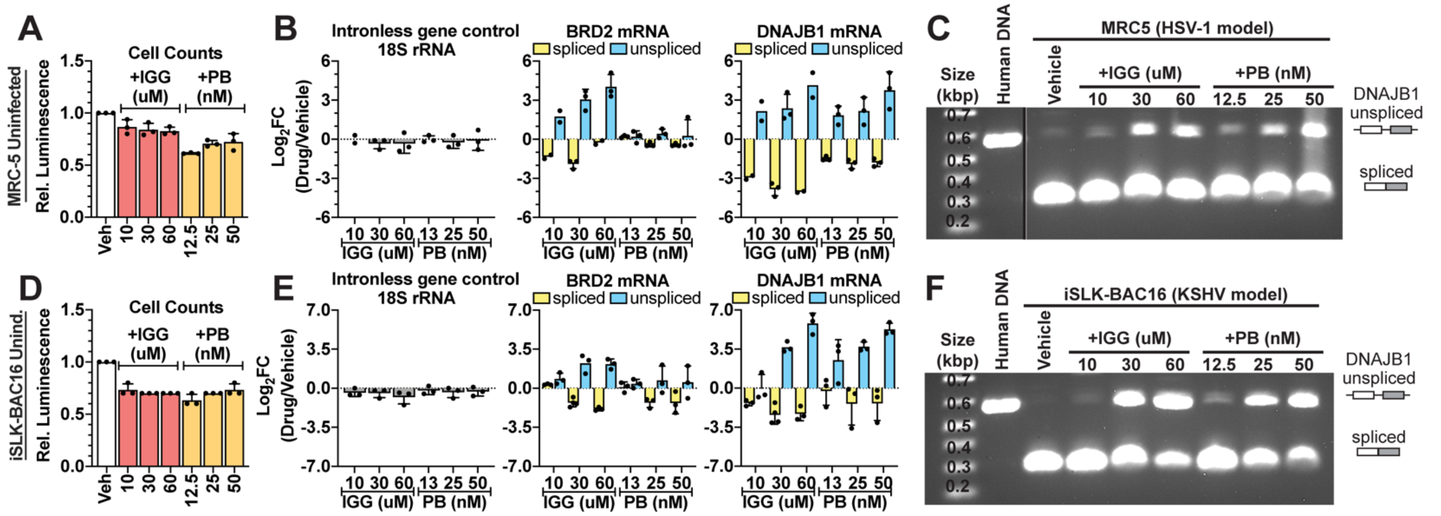


**Supplementary Figure 4-1. Impact of spliceosome inhibition on infection models**

10, 30, 60 uM Isoginkgetin (IGG) or 12.5, 25, 50 nM Pladienolide B (PB) were added for 24 hours to either A-C) uninfected MRC-5 cells or D-F) Uninduced iSLK-BAC16. Data points are biological replicates, column bars are the average, and error bars are standard deviation. A, D) Cell viability was assessed using CellTiterGlo and plotted as luminescence relative to the DMSO or vehicle (Veh) treated control. B, E) To detect spliced mRNA, RNA was reverse transcribed using oligo-dT primers, qPCR primers spanned exon-exon junctions, and relative to cDNA qPCR standard curves. To detect unspliced mRNA, RNA was reverse transcribed using random decamers, qPCR primers within intronic regions, and relative to purified genomic DNA standard curves. Data is plotted as the $\log_2$ fold change (inhibitor/vehicle). C, F) RNA or purified human genomic DNA was reverse transcribed and PCR amplified using primers which bound upstream of an intron within the DNAJB1 gene. PCR product was run on an agarose gel and stained with ethidium bromide.

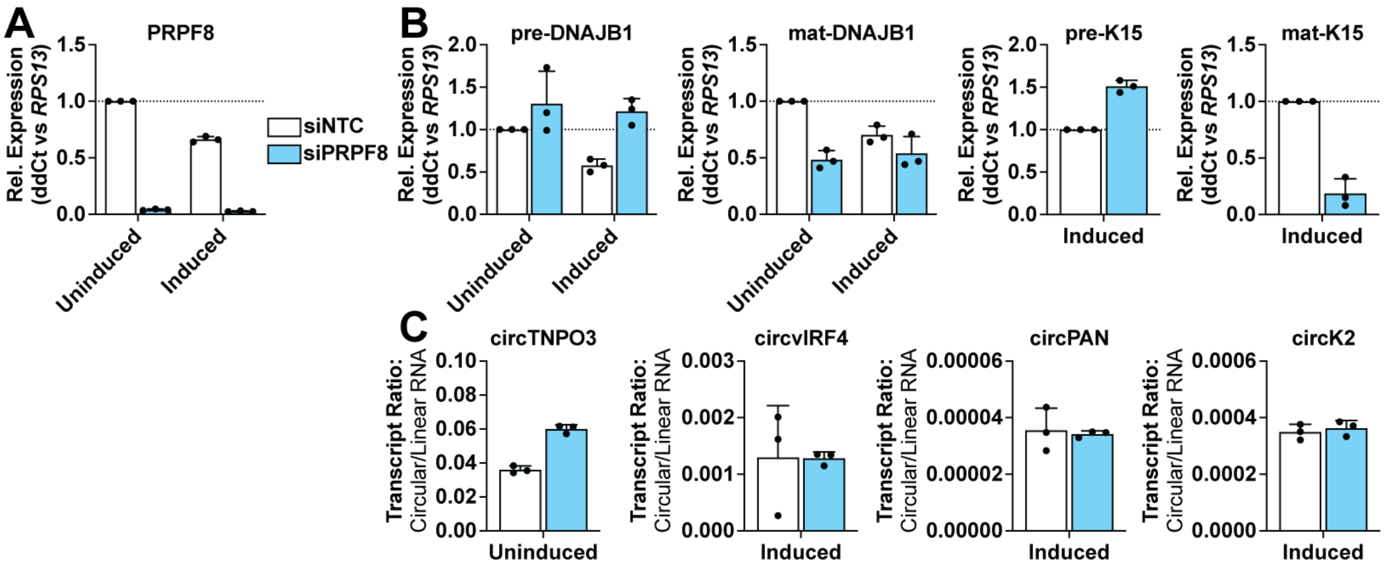


**Supplementary Figure 4-2. Impact of spliceosome depletion on KSHV circRNA levels.**
iSLK-BAC16 were transfected with siRNAs targeting PRPF8 or a nontargeting control (NTC) for 8 hours. Subsequently cells were treated with vehicle (uninduced) or Dox and NaB (induced) for 64 hours. RNA was collected at 72 hours after addition of siRNAs. Transcripts were quantified by A-B) qPCR or C) digital droplet PCR. A-B) qPCR data was analyzed as ddCt relative to the reference gene, *RPS13*. C) Transcript ratios were determined using divergent (circular) and convergent (linear) primers. Data points are biological replicates, column bars are the average, and error bars are standard deviation.

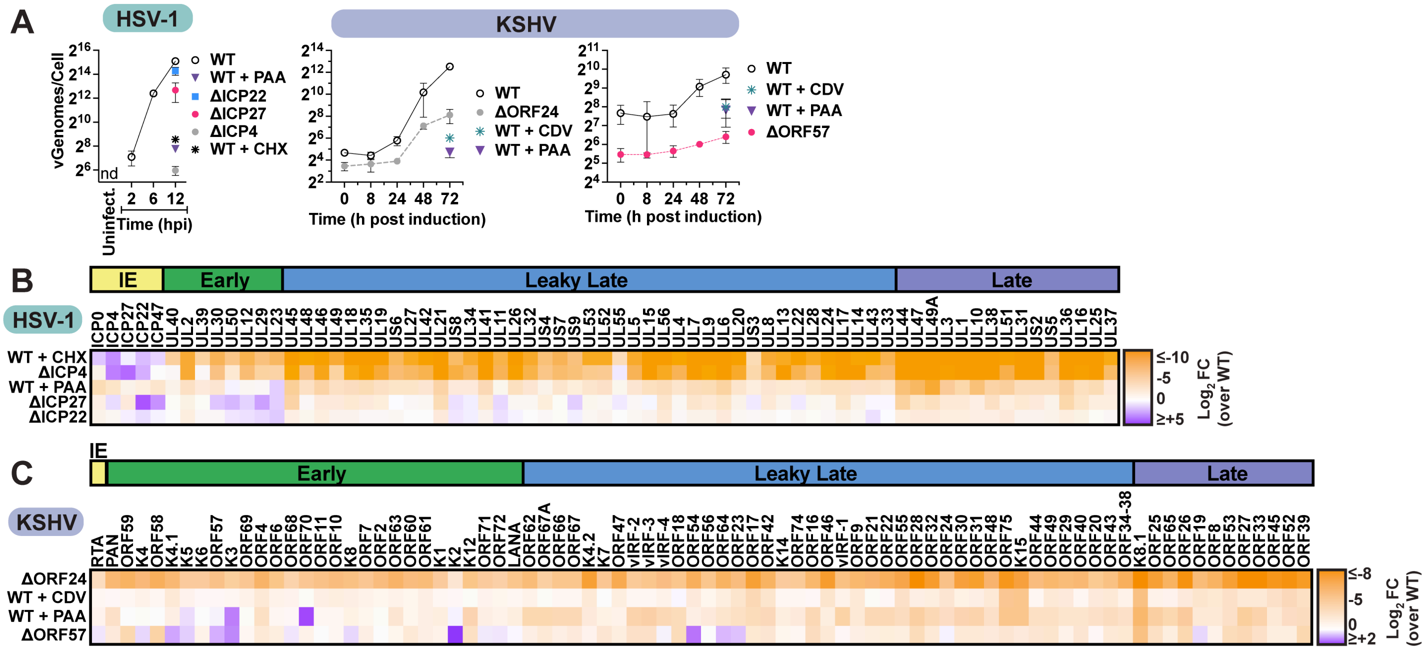

**Supplementary Figure 5-1. HSV-1 and KSHV gene expression in wildtype and perturbed models** A) qPCR assessment of HSV-1 (n=3) or KSHV (n=2) genome quantity, plotted as the number of viral genomes per cell. Data points are the average and error bars are standard deviation. B-C) Viral gene expression for RNA-Seq data in Fig. 5 normalized to ERCC spike-in controls, plotted as $Log_2FC$ relative to a paired wildtype. Data is the average of biological duplicates. Genes are clustered by transcriptional class and labeled as LAT (latent), IE (immediate early), Early, Leaky Late (L1), Late (L2).

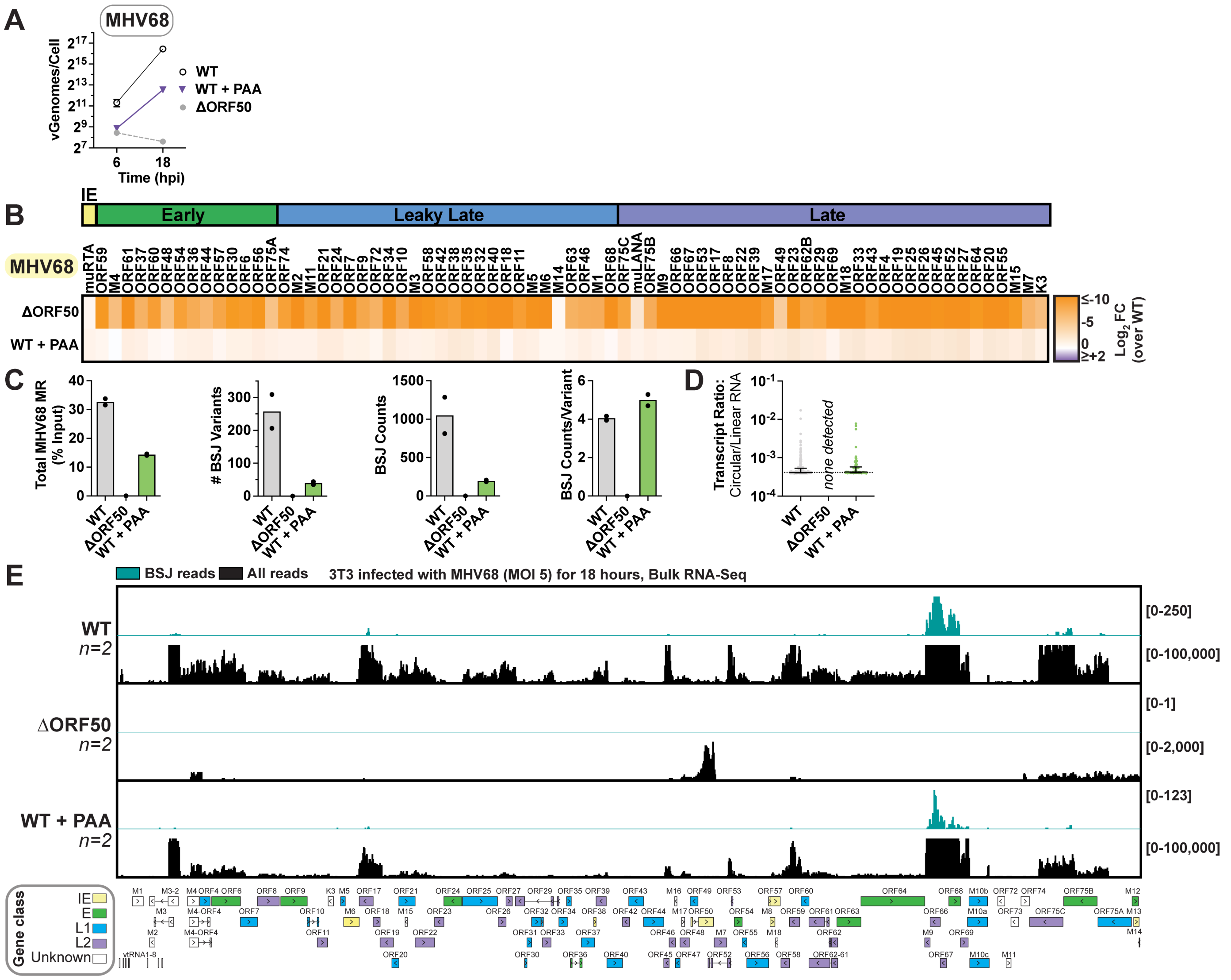


**Supplementary Figure 5-2. MHV68 gene expression in wildtype and perturbed models**

3T3 infected with wildtype MHV68 or ΔORF50 virus at an MOI of 5 PFU/cell. If indicated, 100 ug/mL phosphonoacetic acid (PAA) was added at 1.5 hours post infection. A) qPCR assessment of genome quantity (n=3), plotted as the number of viral genomes per cell. Data points are the average and error bars are standard deviation. B-D) RNA was collected at 18 hours and RNA-Seq performed for biological duplicates. Gene counts were quantified using RNA STAR and normalized to ERCC spike-in reads. BSJ counts and Circular/Linear ratios were quantified using CHARLIE. B) Viral gene expression plotted as $Log_2FC$ relative to a paired wildtype. Data is the average of biological duplicates. Genes are clustered by transcriptional class and labeled as IE (immediate early), Early, Delayed Early, and Late. C) Sequencing overview for all viral mapped reads (MR), as percent total reads (% Total). The number of unique BSJ variants, total BSJ counts, or BSJ counts/variant is reported for high confidence viral circRNAs. Each point is a biological replicate, column bars are the average. D) Circ/Linear ratios for high confidence viral BSJ variants. Each dot represents a unique BSJ, cross-bars are the geometric mean and error bars are the geometric standard deviation. E) Visualization of high confidence circRNAs for MHV68. Blue and black traces include circular (back spliced reads) and linear (non-chimeric) reads, respectively. Traces are the sum of raw BSJ or linear read values for all biological replicates. Y-axis minimum and maximum values are shown on the right. Viral genes are shown below and labeled by gene class as IE (yellow), E (green), L1 (blue), L2 (purple), and unknown (white).

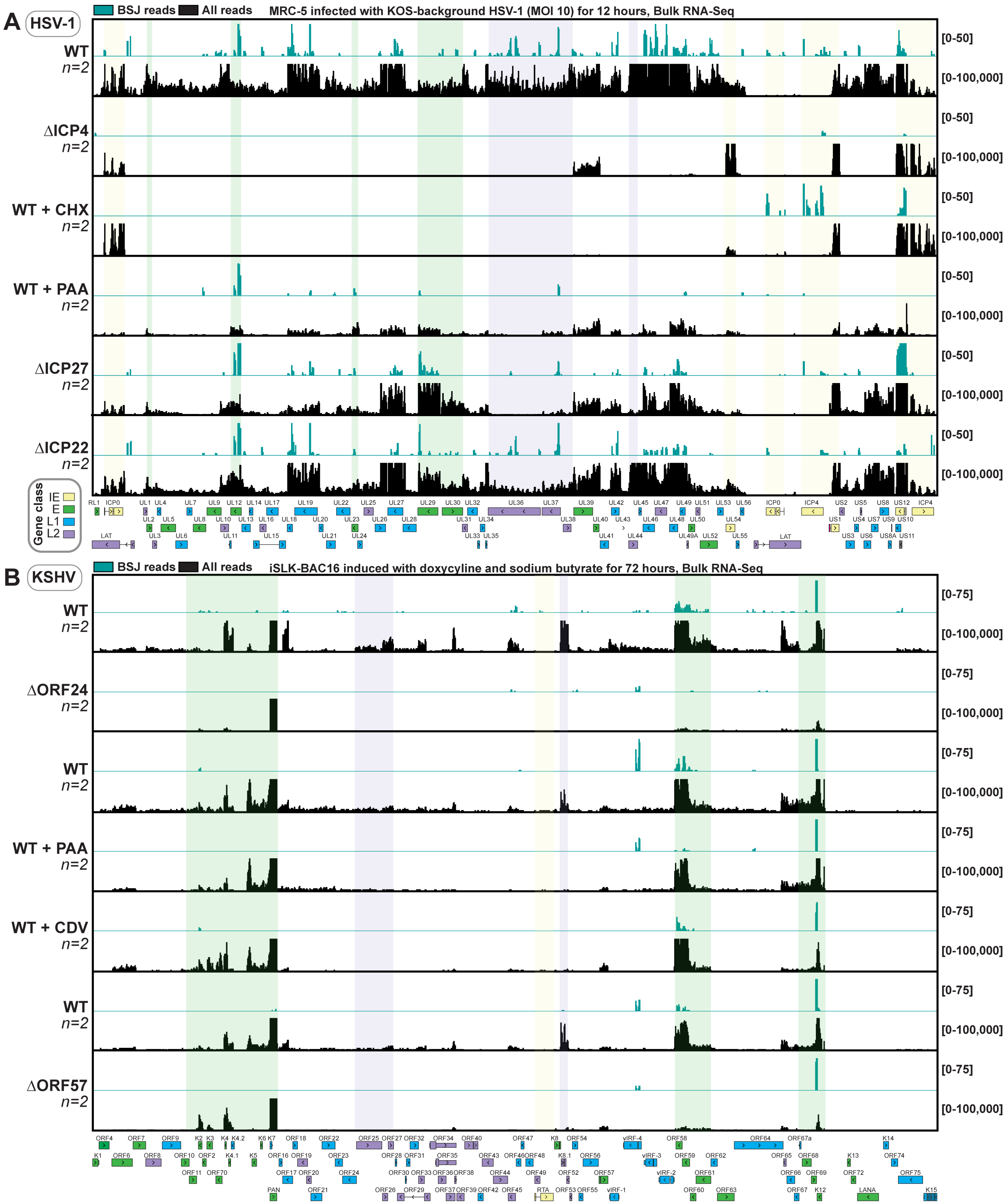


**Supplementary Figure 5-3. HSV-1 and KSHV transcript profiles in wildtype and perturbed models**

Visualization of high confidence HSV-1 and KSHV circRNAs. Blue and black traces include circular (back spliced reads) and linear (non-chimeric) reads. Traces are the sum of raw BSJ or linear read values for biological duplicates. Y-axis minimum and maximum values are shown on the right. Viral genes are shown below and labeled by gene class as IE (yellow), E (green), L1 (blue), and L2 (purple).

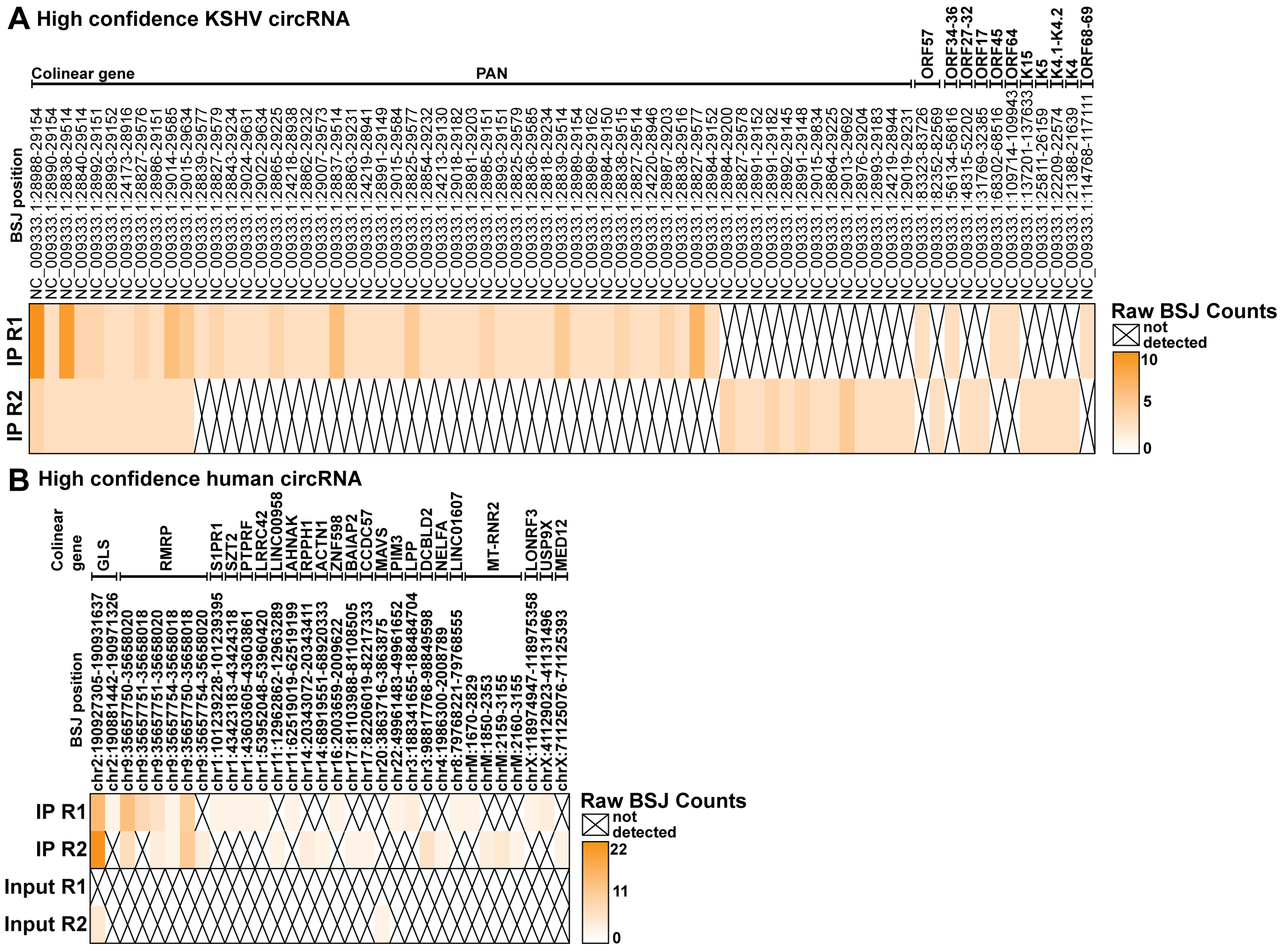


**Supplementary Figure 6-1. ORF57 eCLIP heatmap**

A-B) ORF57 eCLIP (n=2) was performed on iSLK-BAC16 treated with sodium butyrate and doxycycline for 24 hours. “Input” is a paired, size-selected RNA-Seq wherein ORF57 immunoprecipitation was not performed. CircRNA were quantified using CHARLIE and plotted as raw BSJ counts. BSJ position is listed with colinear genes labeled above. Heatmaps include all high confidence A) viral and B) host circRNAs in the dataset.

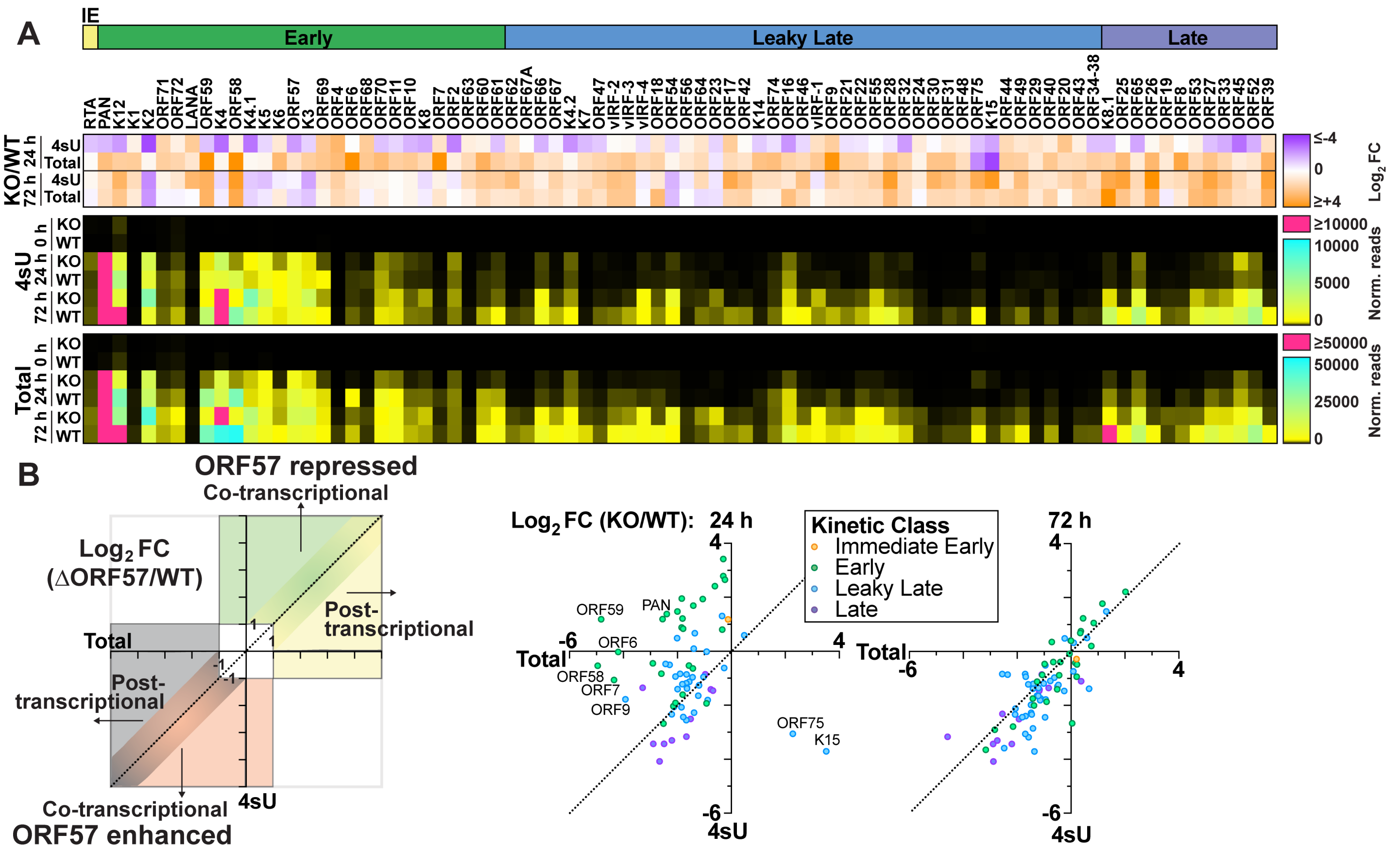

**Supplementary Figure 6-2. ORF57 regulation of non-chimeric viral transcripts**

Bulk (total) and 4-thiouridine (4sU) RNA-Seq for ΔORF57 (KO) and wildtype (WT) iSLK-BAC16 during lytic reactivation. iSLK-BAC16 were treated with (24 and 72 h) or without (0 h) 1 mM sodium butyrate and 1 ug/mL doxycycline. Gene counts were quantified using RNA STAR and normalized for 4sU and total RNA-Seq as Reads Per Kilobase per Million mapped reads (RPKM) or relative to ERCC spike-ins, respectively. Data is the average of biological duplicates. A) Viral gene expression plotted as normalized reads (norm. reads) or ΔORF57 relative to a paired wildtype control ($log_2FC$). Genes are clustered by transcriptional class and labeled as IE (immediate early), Early, Leaky Late, and Late. B) Viral gene expression from 24 or 72 hours post induction plotted as $log_2FC$ (KO/WT), where the x-axis is total RNA-Seq data and the y-axis is 4sU-Seq data.

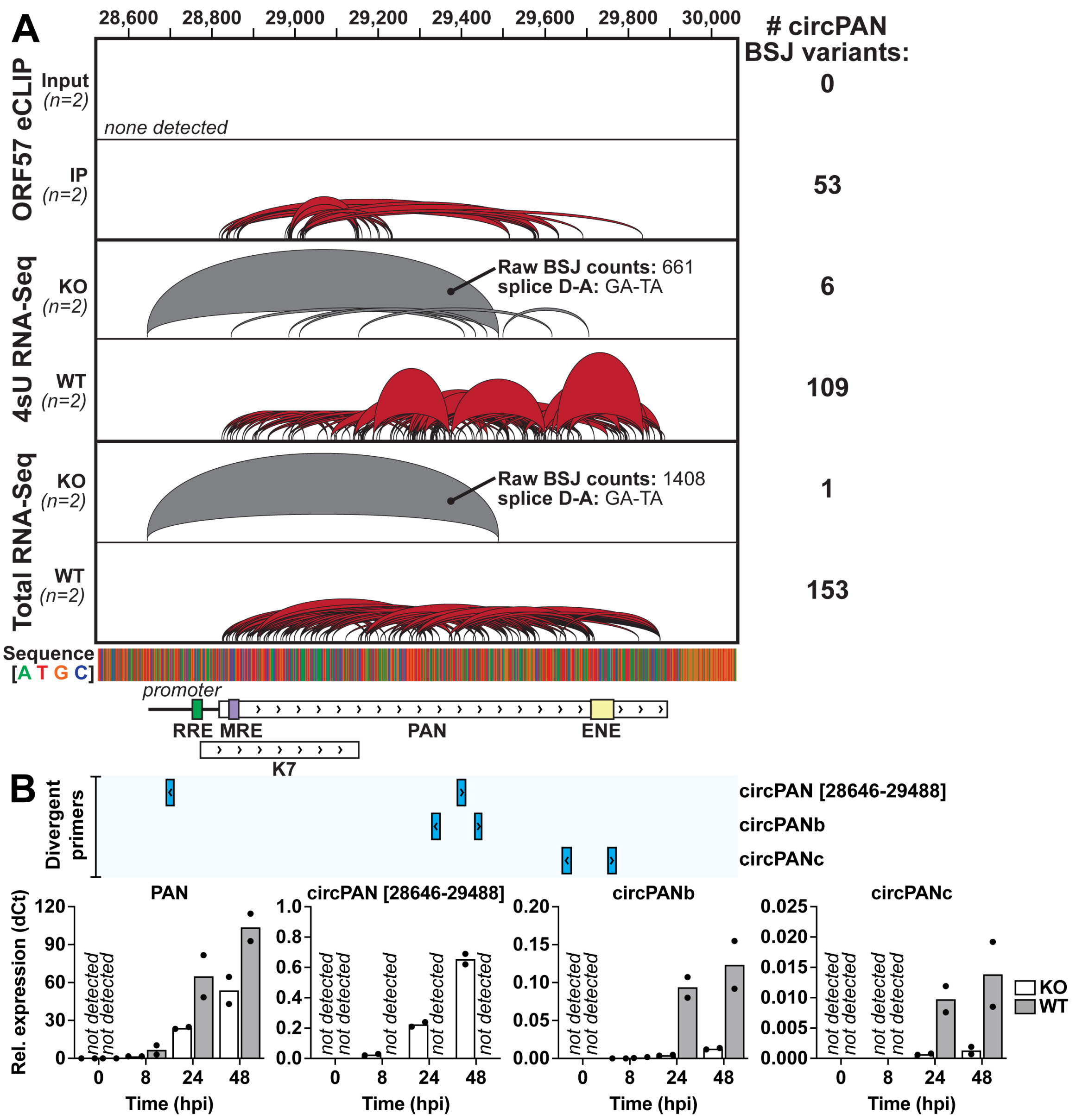


**Supplementary Figure 6-3. ORF57 dependence of PAN BSJ variants**

A) Visualization of high confidence circRNAs at the PAN locus. ORF57 eCLIP (n=2) was performed on induced iSLK-BAC16 (24 h). “Input” is a paired, size-selected RNA-Seq wherein ORF57 immunoprecipitation was not performed. Total and 4sU RNA-Seq (n=2) was performed on ΔORF57 (KO) or iSLK-BAC16 (WT) cells with (24 h) or without (Unind.) reactivation. Sashimi plots show high confidence circRNA with arcs proportional to raw BSJ counts. Each dataset is the sum of biological duplicates. Viral genes and domains including RTA response element (RRE), MTA response element (MRE), expression and nuclear retention element (ENE) are shown below. B) RNA was collected from ΔORF57 (KO) or iSLK-BAC16 (WT) cells reactivated with doxycline and sodium butyrate for 0, 8, 24, and 48 hours. PAN and circPAN transcripts were measured using qPCR and convergent or divergent primers, respectively. The genomic position of divergent primer sets is shown relative to the PAN locus. Data is plotted as relative expression: $2^{-1\times(Experimental\ gene\ Ct - Reference\ gene\ Ct)}$, using 18S rRNA as the reference gene. Individual data points are biological replicates, column bars are the average.
